# Loss of Parkin Disrupts Nuclear and Mitochondrial Programs Required for Muscle Regeneration

**DOI:** 10.64898/2026.03.20.712989

**Authors:** Melissa Gourlay, Mah Rukh Abbasi, George Cairns, Madhavee Thumiah-Mootoo, Jeremy Racine, Ha My Ly, Anna Wang, Nikita Larionov, Alexandre Blais, Mireille Khacho, Yan Burelle

## Abstract

Skeletal muscle stem cells (MuSCs) rely on precisely coordinated metabolic and nuclear transitions to exit quiescence, enter the cell cycle, and regenerate tissue. How these processes are coupled remains poorly defined. Here, we identify PARKIN as a critical integrator of mitochondrial quality control and nuclear RNA processing programs that together enable balanced MuSC lineage progression. Using a MuSC-specific, inducible Park2 knockout model, we show that PARKIN supports mitophagy in quiescent MuSCs, and its loss triggers premature mitochondrial polarization and fragmentation — hallmarks of metabolic activation — that compromise appropriate self-renewal and fate specification. Unexpectedly, MuSCs harbor a constitutive nuclear pool of PARKIN that rises rapidly upon activation and localizes to interchromatin regions, with focal association with nuclear speckles. Park2-deficient MuSCs exhibit transcriptomic signatures consistent with widespread RNA isoform switching and intron retention, particularly affecting splicing machinery components, accompanied by altered nuclear speckle organization and impaired cell cycle progression. These findings reveal that PARKIN safeguards both mitochondrial homeostasis and the RNA processing architecture essential for activation, thereby coordinating metabolic and nuclear reprogramming during early MuSC state transitions. Our work positions PARKIN as a dual compartment regulator required for robust skeletal muscle regeneration.

## 1 INTRODUCTION

Skeletal muscle exhibits remarkable regenerative capacity, which is primarily driven by muscle stem cells (MuSCs), a population of tissue-resident stem cells located in a specialized niche beneath the basal lamina of myofibers^1–4^. In homeostatic conditions, MuSCs persist in a reversible state of cell cycle arrest known as quiescence. Following injury, these cells are rapidly activated, re-enter the cell cycle, and expand through proliferative divisions. The majority of activated MuSCs commit to the myogenic lineage to support tissue repair, while a subset retains stem cell identity to maintain the quiescent pool ^1–4^. Successful regeneration is thus contingent on the coordination of MuSC activation, controlled proliferation, and fate specification, ensuring a proper balance between self-renewal and differentiation ^1–4^. Although these processes are critical for preserving long-term regenerative potential, the molecular networks that integrate these processes remain incompletely defined.

Regulation of MuSC behavior was initially viewed primarily through the lens of transcription factor networks and signaling pathways that regulate cell cycle re-entry and orchestrate the balance between self-renewal and commitment to the myogenic lineage. This framework emphasized gene regulatory circuits and extrinsic niche-derived cues as the dominant forces guiding stem cell behavior. However, emerging evidence has revealed additional complexity by revealing that MuSC metabolism is an active regulator of fate decisions ^5–10^. Mitochondria, in particular, have gained recognition as central hubs that integrate bioenergetic demands with signaling and epigenetic control^5,8–10^. Changes in mitochondrial function—through shifts in oxidative phosphorylation (OXPHOS), reactive oxygen species (ROS) levels, or the availability of key metabolites such as NAD⁺, acetyl-CoA, or α-ketoglutarate—have been shown to directly influence chromatin accessibility, transcriptional programs, and post-translational modifications ^5–9,11^. These discoveries have redefined our understanding of MuSC regulation, positioning mitochondrial and metabolic state as dynamic and instructive elements in stem cell fate determination.

Maintaining mitochondrial quality and promoting specific functional properties, including appropriate levels of oxidative phosphorylation, membrane potential, and reactive oxygen species, is considered essential not only to preserve the quiescent state but also to enable efficient transitions into activation and proliferation ^5,6,12,13^. These mitochondrial properties differ between quiescent and activated MuSCs and must be coordinated with nuclear programs that define stem cell state, including transcriptional reprogramming, epigenetic remodeling, modulation of mRNA processing, and translation ^14–17^. Transitions out of quiescence require silencing of quiescence-enforcing gene networks, activation of cell cycle regulators, and increased biosynthetic capacity ^14–17^ However, mechanisms that coordinate mitochondrial state with these nuclear programs remain poorly defined.

One candidate coordinator is the E3 ubiquitin ligase PARKIN, which has established roles in mitochondrial quality control and emerging links to nuclear processes^5,18–20^. In addition to its canonical role in mitochondrial surveillance, PARKIN has been implicated in the regulation of metabolic homeostasis and cell cycle progression ^21–24^, positioning it as a potential integrator of metabolic status with cellular state. While PARKIN has been extensively studied in post-mitotic tissues such as neurons, its function in MuSCs, where it is expressed^13^ remains ill-defined. In this study, we investigated the role of PARKIN in regulating MuSC function. We show that PARKIN actively regulates mitophagy and mitochondrial properties during quiescence and transition to activation, ensuring a proper balance between self-renewal and commitment. Unexpectedly, we identify a distinct nuclear pool of PARKIN and find that PARKIN is required for proper nuclear speckle organization, splicing fidelity, and cell cycle progression. Our findings suggest that PARKIN coordinates both mitochondrial and nuclear programs essential for productive MuSC activation and expansion.

## 2 RESULTS

### 2.1 Mitophagy marks quiescence and rapidly fades with activation

Mitophagy dynamics was first characterized during the transition from quiescence to activation. Given that MuSCs are exquisitely sensitive to niche perturbations, with enzymatic dissociation and FACS purification alone being sufficient to trigger early activation events ^15,25,26^, we assessed mitophagy readouts across three experimental conditions that capture different cellular states: MuSCs from *in situ* PFA-fixed uninjured muscle (fixed before isolation) corresponding to the native quiescent state, MuSCs fixed immediately after isolation corresponding to the quasi-quiescent/early activation phase induced by isolation procedures, and MuSCs fixed after 4 hours of culture corresponding to further activation progression.

High-resolution confocal microscopy and three-dimensional reconstruction of TOM20-labeled mitochondria and LC3-labeled autophagosomes was first performed to obtain a global readout of mitophagy. These experiments revealed a strikingly high level of colocalization between mitochondria and autophagosomes, in *in situ* fixed quiescent MuSCs, representing more than 20% of cellular mitochondrial content (Fig. 1A-C). This colocalization then declined progressively in post-isolation fixed and 4-hour cultured cells (Fig. 1A-C) and was unrelated to changes in mitochondrial biomass and/or autophagosome content (Fig. S1A-B), pointing to rapid changes in mitophagy dynamics during the quiescence-to-activation phase. Previous work using this approach has established that such dynamics reflect reduced mitophagy flux rather than increased lysosomal degradation ^13^, and we confirmed lysosomal activity remains negligible in MuSCs at these early time points using MuSCs from the EGFP-RFP-LC3 reporter mouse line (Fig. S1C-D).

**Figure 1:**
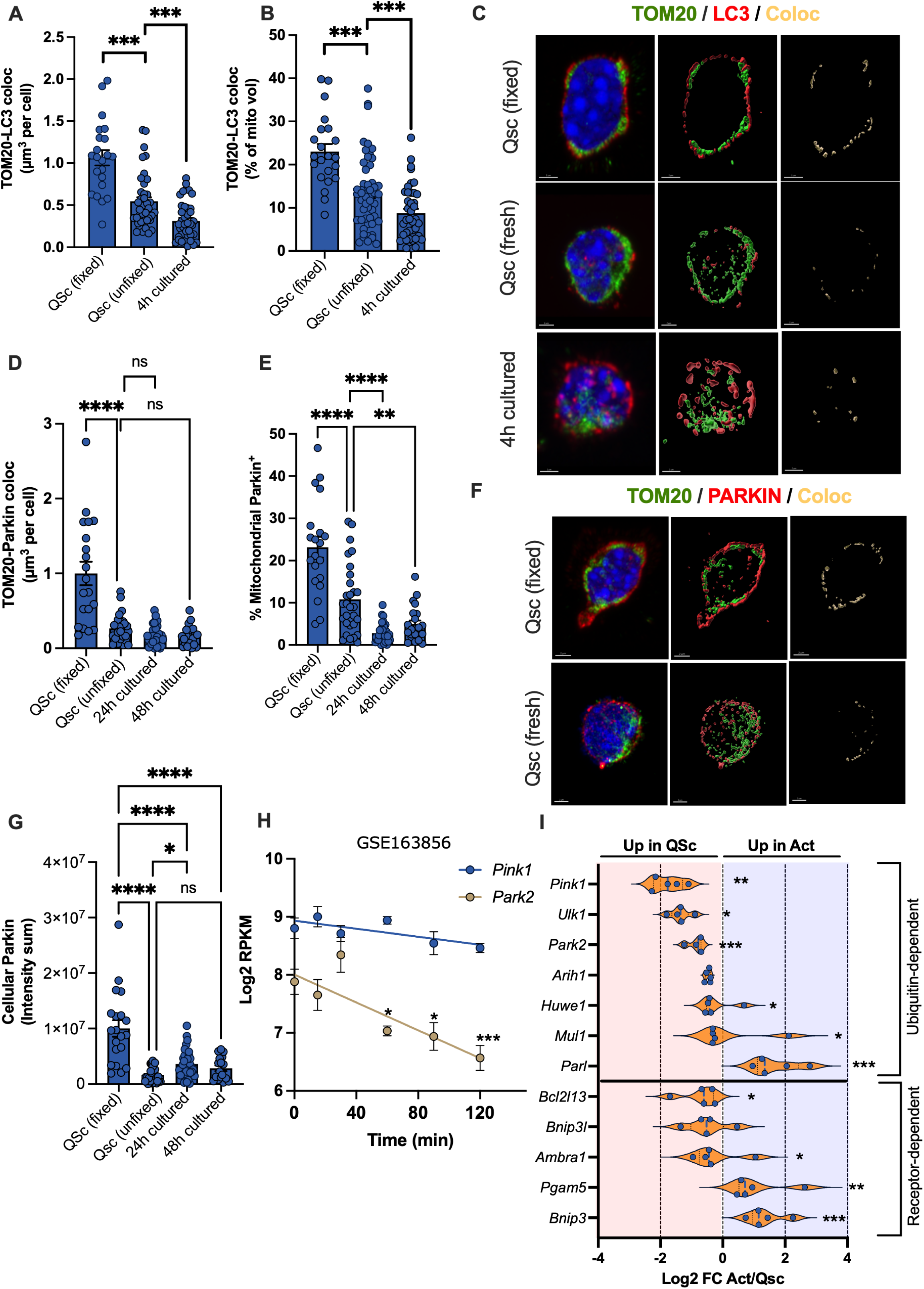
Mitophagy marks quiescence and rapidly fades with activation. **A-B**) Volume of mitochondria colocalized to autophagosomes expressed in absolute value or relative to total mitochondrial volume in *in situ* fixed, freshly sorted quiescent and 4h *in vitro* activated MuSCs (*n*=48-55 cells from 3 mice in each group and experimental condition). **C)** Confocal image and 3D reconstruction of TOM20-labeled mitochondria (green) and LC3-labeled autophagosomes (red) in each of the experimental conditions examined. The volume of mitochondria overlapping with autophagosomes is shown in yellow in the right-end panels, where mitochondria and autophagosomes surfaces have been removed to highlight changes in colocalization. **D)** Colocalization of PARKIN to TOM20-labeled mitochondria expressed in absolute volume of PARKIN^+^ structures overlapping with mitochondria per cell (*n*=25-36 cells from 3 mice in each group and experimental condition). **E)** Proportion of TOM20-labeled mitochondria labeled with PARKIN, computed using data presented in panel D and total mitochondrial content (Fig. S1A). **F)** Confocal image and 3D reconstruction of TOM20-labeled mitochondria (green) and PARKIN-labeled structures (red) in each of the experimental conditions examined. The volume of PARKIN overlapping with mitochondria is shown in yellow in the right-end panels, where mitochondria and PARKIN^+^ surfaces have been removed to highlight changes in colocalization. **G)** Total cellular PARKIN content in the indicated experimental conditions. **H)** *Parkin* transcript levels in MuSCs purified from muscles that were fixed *in situ* in healthy uninjured conditions or at 15, 30, 60, 90 and 120 min after CTX injury. Data is taken from (GSE163856 ^26^). All data are presented as mean ± SEM. ns: not significant, *: p<0.05, **: p<0.01, ***: p<0.001, ****: p<0001 on unpaired two-tailed *t* tests or one-way ANOVAs. **I)** Changes in the transcript levels of ubiquitin- and receptor-dependent mitophagy in three transcriptomics datasets (GSE70736, GSE55490, GSE47177 ^12,17,27^) that compared quiescent and *in vivo* activated MuSCs 36-72h following muscle injury with CTX or BaCl_2_. Each dot represents data from an individual dataset and a specific timepoints. Genes shown where differentially expressed in all datasets with a q value of less than 0.05. Level of statistical significance shown is based on the average q value.

Several analyses were then performed to characterize the involvement of PARKIN. Confocal microscopy and 3D reconstruction revealed that the volume of PARKIN-positive signal in whole cells and mitochondria was highest in *in situ* fixed quiescent MuSCs and declined upon gradual activation (Fig. 1D-F), mirroring changes in mitochondrial colocalization to autophagolysosomes (Fig. 1A-C). Consistent with reduced PARKIN levels, analysis of bulk RNAseq data comparing MuSCs that were fixed at early times points (0-120 min^25,26^) following *in vivo* cardiotoxin (CTX) injury also revealed high levels of *Parkin* transcripts in quiescent cells followed by a rapid decline in response to progressive activation, with *Pink1* showing a similar but non-significant decline (Fig. 1H). Broader analysis of transcriptomics datasets comparing unfixed MuSCs purified from uninjured and injured muscles 36–72 h post-injection of CTX or BaCl_2_ ^12,17,27^ also revealed a systematic enrichment of *Pink1* and *Parkin* transcripts in the quiescent compared to the activated state (Fig. 1I). Together, these results indicate that PARKIN expression is highest in quiescence and declines during activation, suggesting that PARKIN is actively involved in the maintenance of MuSC quiescence and rapid transition to the activation state.

### 2.2 Parkin regulates mitophagy from quiescence to activation

To investigate PARKIN’s role in regulating MuSC function, *Parkin* was selectively inactivated in MuSCs using *Pax7CreERT2*-mediated recombination. Successful *Parkin* inactivation was confirmed by qPCR and immunofluorescence imaging of unfixed MuSCs freshly isolated from uninjured muscle one week after the final tamoxifen treatment (Fig. 2A-C). To probe for potential compensatory effects, transcript levels of key regulators of ubiquitin- and receptor-dependent mitophagy were examined and found to be comparable between control and PARKIN-deficient MuSCs, indicating no transcriptional compensation secondary to the loss of *Parkin* (Fig. 2D-E). Consistent with a role for PARKIN in regulating mitophagy, colocalization between mitochondria and autophagosomes was reduced by 30-40% particularly in quiescent, but also in the activated state (Fig. 2F-H), in line with the reduced colocalization and mitophagy flux we previously reported in MuSCs from germline *Pink1^-/-^* mice ^13^.

**Figure 2:**
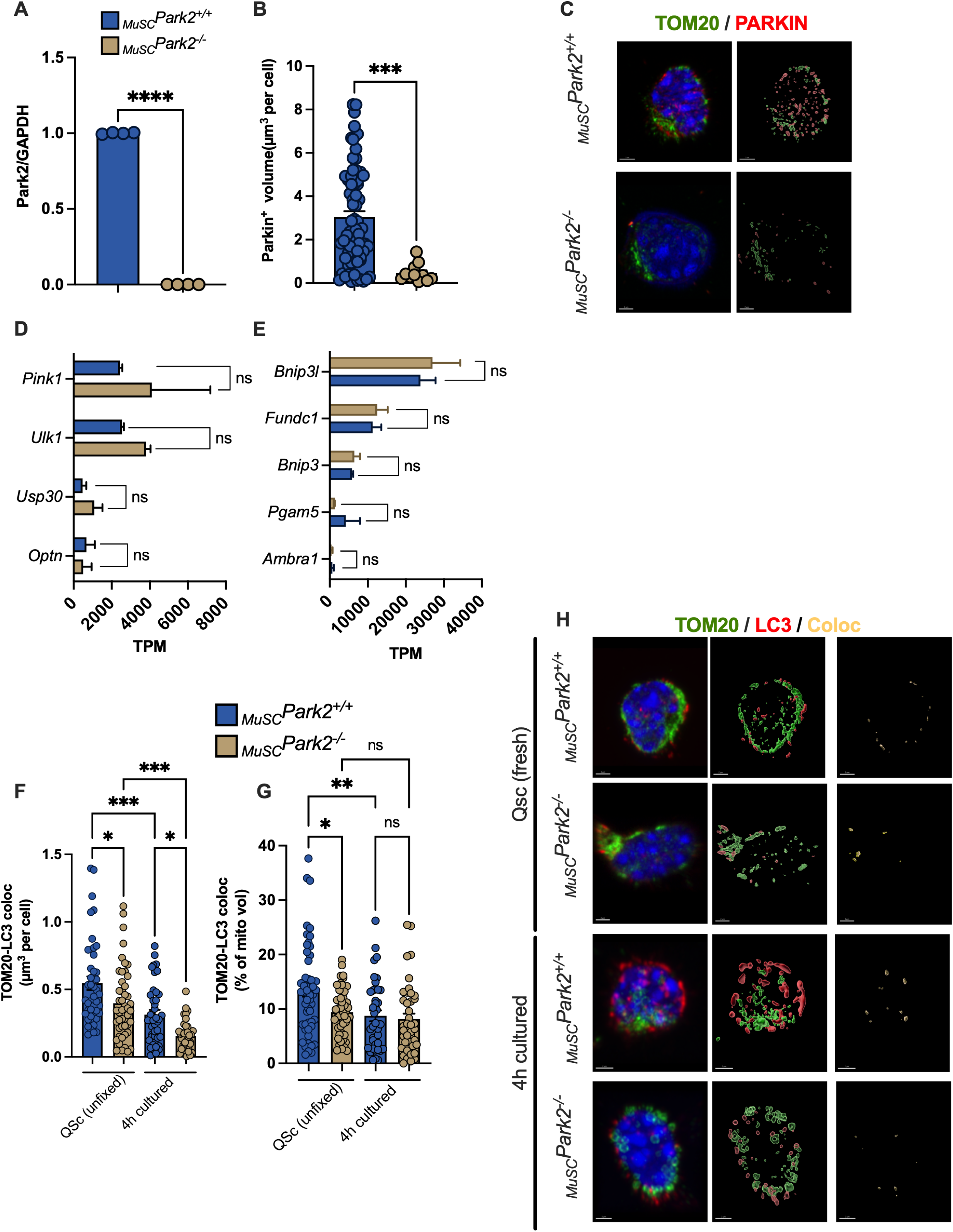
PARKIN regulates mitophagy from quiescence to activation. **A-C)** Parkin transcript and protein levels in FACS-purified MuSCs from *_MuSC_Park2^+/+^* and *_MuSC_Park2^-/-^* mice 1 week after the last tamoxifen treatment (*n*=4 mice per group). Protein abundance was quantified as the total volume of PARKIN^+^ structures (B) in cells labeled with antibodies against PARKIN and TOM20 (*n*=11-70 cells from 2-4 mice per group,) (C). **D-E)** transcript abundance expressed in Transcript Per Million (TPM) for genes involved in Parkin- and Receptor-mediated mitophagy (*n*=3 mice per group). **F-G)** Volume of mitochondria colocalized to autophagosomes expressed in absolute value or relative to total mitochondrial volume in freshly sorted quiescent and 4h *in vitro* activated MuSCs (*n*=41-52 cells from 3 mice in each group). **H)** Confocal image and 3D reconstruction of TOM20-labeled mitochondria (green) and LC3-labeled autophagosomes (red) in each of the experimental conditions examined. The volume of mitochondria overlapping with autophagosomes is shown in yellow in the right-end panels, where mitochondria and autophagosomes surfaces have been removed to highlight changes in colocalization. All data are presented as mean ±SEM. ns: not significant, *: p<0.05, **: p<0.01, ***: p<0.001, ****: p<0001 on unpaired two-tailed *t* tests or one-way ANOVAs.

### 2.3 Parkin loss disrupts regeneration and skews fate toward commitment over self-renewal

To examine the functional impact of PARKIN deficiency on MuSC biology, muscle injury was induced through unilateral CTX injection in the *Tibialis Anterior*, then regeneration dynamics and MuSC population changes were monitored over time. *_MuSC_Park2^-/-^* mice exhibited compromised muscle regeneration, as evidenced by persistently reduced fiber cross-sectional area from 14 days post-injury onward compared to controls (Fig. 3A-C). PAX7^+^ MuSC numbers were also consistently diminished in PARKIN-deficient mice, with reductions evident between 4-14 days post-injury suggesting defective pool expansion and/or compromised fate specification, and sustained decreases at 21 days suggesting incomplete restoration of the quiescent MuSC reservoir (Fig. 3D-E).

**Figure 3:**
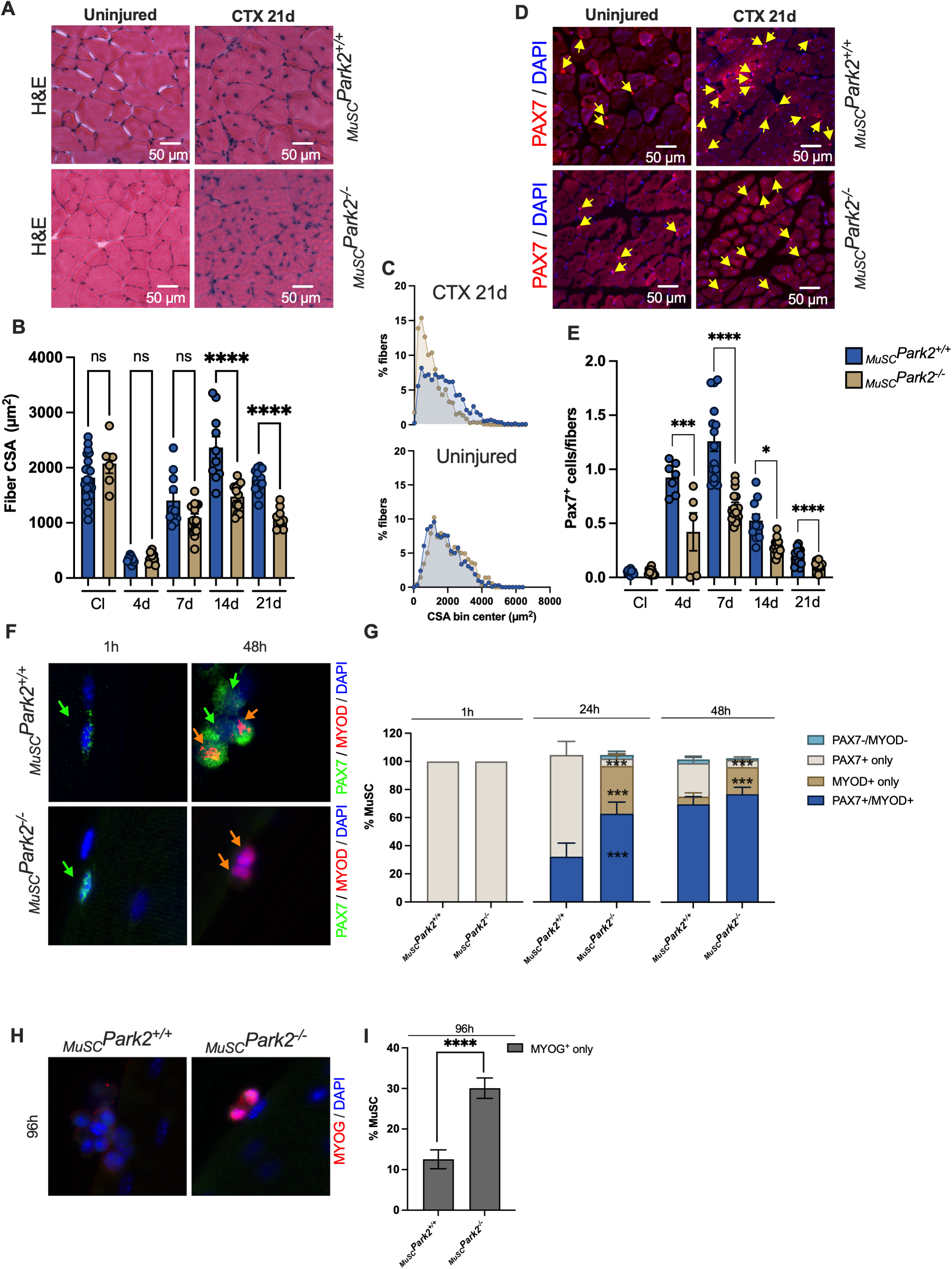
PARKIN loss disrupts regeneration and skews fate toward commitment over self-renewal. **A)** Representative H&E stains of fibers in contralateral uninjured and injured TA muscle of *_MuSC_Park2^+/+^*and *_MuSC_Park2^-/-^* mice 21 days after CTX injury. **B)** Mean cross sectional area (CSA) of fibers in contralateral (Cl) uninjured and injured TA muscle at 4, 7 14- and 21-days post CTX injury (*n*=6-21 ROIs from 3-5 TA muscle per group). **C)** Fiber size distribution of uninjured and injured muscles at 21 days post injury. **D)** Representative images of PAX7^+^ and DAPI stained TA cross sections in uninjured and injured TA 21 days post CTX injury. **E)** Number of PAX7^+^ positive cells in uninjured and injured TA muscles at 4, 7 14 and 21-days post CTX injury. Given the differences observed in fiber CSA between genotypes, cell count is expressed as the number of PAX7^+^ cells divided by the number of fibers present in each 500 µm^2^ ROI analyzed (*n*=5-15 ROIs from 3-5 TA muscle per group). **F)** Representative images of PAX7 and MYOD labeled MuSCs at the surface of EDL fibers after 1 and 48h of culture. **G)** Proportion of quiescent (PAX7^+^ only) and activated MuSCs (PAX7^+^/MYOD^+^) and MYOD^+^ only at 1, 24 and 48 hours post isolation (*n*=65-78 cells from 13-16 fibers isolated from 3-4 mice per genotype). **H)** Representative images of PAX7 and MYOG labeled MuSCs at the surface of EDL fibers after 96h of culture. **I)** Proportion of MuSCs expressing MYOG after 96h in culture (n=634-712 MuSCs from 4-5 fibers isolated from 3 mice per genotype). All data are presented as mean ± SEM. ns: not significant, *: p<0.05, **: p<0.01, ***: p<0.001, ****: p<0001 on unpaired two-tailed *t* tests or one-way ANOVAs.

This abnormal regenerative response was further dissected by monitoring MuSC fate decision in cultured EDL myofibers. PARKIN-deficient MuSCs showed an increased propensity for activation and commitment at the expense of self-renewal, with elevated PAX7^+^/MYOD^+^ and PAX7^-^/MYOD^+^ populations and reduced PAX7^+^-only cells at 24 and 48h (Fig. 3F-G). The proportion of cells expressing MYOG^+^ was also increased in absence of PARKIN suggesting that the imbalance in fate decision pushed cells toward precocious differentiation at the expense of self-renewal (Fig. 3H), consistent with the incomplete restoration of the MuSC pool following *in vivo* injury (Fig. 3D-E).

### 2.4 Parkin shapes the mitochondrial phenotype of MuSCs

Previous studies have demonstrated that alterations in mitochondrial functional and structural properties are critical determinants of MuSC fate specification ^13,28,29^. To investigate whether PARKIN deficiency induced mitochondrial changes characteristic of activation-prone state, we assessed key mitochondrial parameters in control and knockout MuSCs. Analysis of cellular ATP content in the absence and presence of oligomycin, an ATP synthase inhibitor, revealed comparable OXPHOS contributions to the total ATP pool between genotypes, indicating that PARKIN deficiency did not fundamentally compromise mitochondrial bioenergetic capacity (Fig. 4A-B). However, PARKIN-deficient MuSCs displayed elevated mitochondrial membrane potential (Fig. 4C), and increased mitochondrial network fragmentation (Fig. 4D), two changes that have all been linked to quiescence exit and activation through increased ROS signaling ^28,30,31^.

**Figure 4:**
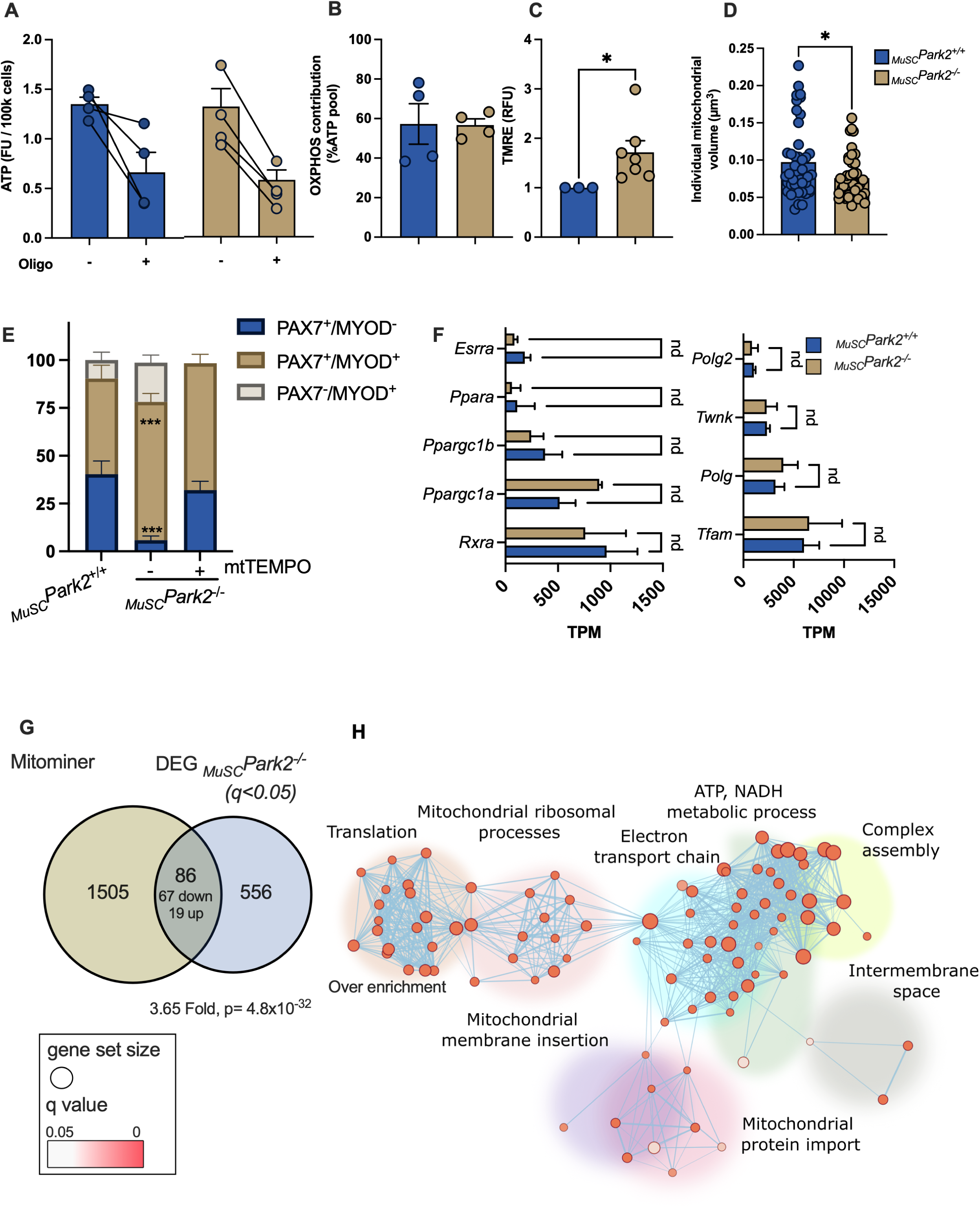
PARKIN shapes the mitochondrial phenotype of MuSCs. **A)** ATP levels in freshly isolated MuSCs from *_MuSC_Park2^+/+^* and *_MuSC_Park2^-/-^*mice measured in absence and presence of the ATP synthase inhibitor Oligomycin (20 µM). **B)** Contribution of OXPHOS to cellular ATP levels calculated as the difference in ATP levels in absence and presence of Oligomycin (*n*=4 MuSC sorts per genotype). **C)** Mean TMRE fluorescence intensity in freshly isolated MuSCs expressed as fold change of control(*n*=3-4 MuSC sorts per genotype). **D)** Average volume of individual mitochondria per cell in freshly isolated MuSCs labeled with TOM20 (n=48-52 cells from from 3 mice in each group). Data was obtained following confocal imaging and 3D reconstruction. **E)** Proportion of quiescent (PAX7^+^ only) activated (PAX7^+^/MYOD^+^) and committed (MYOD^+^ only) MuSCs in EDL fibers cultured in absence or presence of 10 µM mitoTEMPO for 24h (n=52-135 MuSCs from 5 fibers from 4 mice per genotype). **F)** Transcript levels expressed in Transcript Per Million (TPM) for key genes regulating mitochondrial biogenesis and mtDNA replication in freshly isolated MuSCs (n=3 mice per genotype). All data are presented as mean ± sem. nd: not different, *: p<0.05, **: p<0.01, ***: p<0.001, ****: p<0001 on unpaired two-tailed *t* tests or one-way ANOVAs comparing *_MuSC_Park2^+/+^*and *_MuSC_Park2^-/-^*. **G)** Venn Diagrams presenting the number of mitochondrial genes (MitoMiner mitochondrial localisation database^59^) that are differentially enriched in freshly isolated PARKIN deficient MuSCs compared to controls. Fold enrichment of mitochondrial genes is shown below Venn diagram along with the hypergeometric test p value. **H)** g:Profiler enrichment map illustrating the main mitochondrial processes downregulated in PARKIN deficient MuSCs compared to controls. The auto-annotation tool was used on Cytoscape to automatically generate cluster labels.

EDL myofibers were cultured in the presence of MitoTEMPO, a mitochondria-targeted superoxide scavenger (Fig. 4E). MitoTEMPO treatment significantly reduced the proportion of committed (PAX7-/MYOD+) cells in PARKIN-deficient cultures and restored the frequency of unactivated (PAX7+/MYOD-) cells to levels approaching those observed in untreated wild-type controls, pointing to mitochondrial ROS production being a mechanistic driver of dysregulated fate decisions in PARKIN-deficient MuSC

Bulk RNAseq data from unfixed MuSCs freshly isolated from uninjured muscle was also interrogated to determine whether at the gene level PARKIN deficiency was associated with transcriptional signatures of mitochondrial dysfunction. Transcripts for key transcription factors and co-activators involved in mitochondrial biogenesis, as well as proteins required for mtDNA replication, were normally expressed in the absence of PARKIN (Fig. 4F). However, among the 642 genes differentially expressed (padj < 0.05) in PARKIN-deficient MuSCs, 86 encoded mitochondrial proteins, pointing to downstream abnormalities (Fig. 4G). The vast majority (67) of these genes were downregulated in PARKIN-deficient cells. Over-representation analysis revealed that these downregulated mitochondrial genes were enriched for functions including mitochondrial translation, protein import and membrane insertion, respiratory complex assembly, and OXPHOS (Fig. 4H and S2A). Complementary gene set enrichment analysis of the full ranked gene list confirmed suppression of OXPHOS and mitochondrial respiratory chain pathways in PARKIN-deficient MuSCs (Fig. S2B-D), a known transcriptomics hallmark of metabolic departure from quiescence ^15,25^.

Overall, these results suggested that PARKIN deficiency disrupts transcriptional programs essential for oxidative metabolism while promoting mitochondrial changes that predispose MuSCs to activation.

### 2.5 A nuclear pool of PARKIN exists in MuSCs and PARKIN loss disrupts nuclear organization and proliferation

Transcriptomic analysis also revealed that PARKIN deficiency disrupted gene expression related to multiple nuclear processes beyond metabolism. Cell cycle-related pathways were extensively downregulated, including G2/M transition and CDK activity regulation, chromosome architecture and centromeric chromatin assembly, kinetochore assembly, spindle formation and chromosome attachment, and chromosome alignment and segregation. RNA processing pathways were also affected, with significant downregulation of precatalytic spliceosome components (Fig. 5A, Fig. S2 B, E, F). This extensive downregulation of nuclear functions, from chromosome architecture to mitotic and splicing machinery, prompting examination of PARKIN subcellular localization.

**Figure 5:**
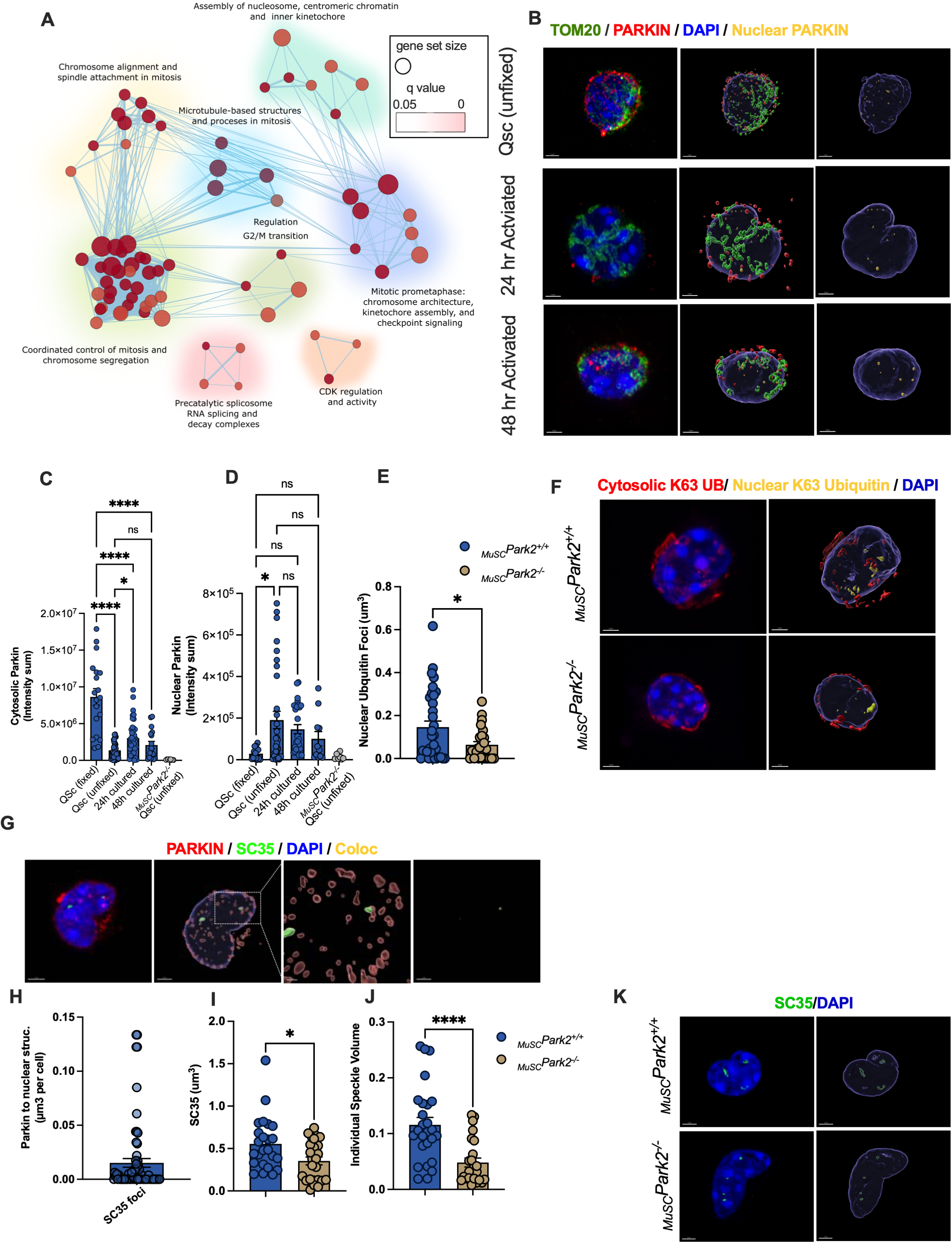
Nuclear PARKIN regulates key processes driving MuSC activation and expansion. **A)** g:Profiler enrichment map illustrating nuclear processes downregulated in PARKIN deficient MuSCs compared to controls. The auto-annotation tool was used on Cytoscape to automatically generate cluster labels. **B)** Confocal image and 3D reconstruction of mitochondria (TOM20, green), nucleus (DAPI, blue) and PARKIN^+^ foci (red) in freshly isolated and 24-48h *in vitro* activated MuSCs from wild type mice. PARKIN^+^ foci located within the nuclear compartment is shown in Yellow in the right-end panels, where mitochondria and nuclear surfaces have been removed for clarity. **C)** Quantification of cytosolic PARKIN^+^ foci in *in situ* fixed, freshly sorted quiescent and 24-48h *in vitro* activated MuSCs. PARKIN was defined as cytosolic if the overlap ratio between PARKIN^+^ foci and nuclear volume ranged between 0 and 0.4. **D)** Quantification of nuclear PARKIN^+^ foci in *in situ* fixed, freshly sorted quiescent and 24-48h *in vitro* activated MuSCs. PARKIN was defined as nuclear if the overlap ratio between PARKIN^+^ foci and nuclear volume was above 0.8 (*n*=10-36 cells from 4 mice per genotype and time point). In panel C and D, note the absence of PARKIN staining in *_MuSC_Park2^-/-^* cells confirming the specificity of the staining. **E)** Quantification of Ubiquitin^+^ foci within the nuclear compartment in *_MuSC_Park2^+/+^* and *_MuSC_Park2^-/-^*MuSCs at 18hr into culture. (*n*=24-32 cells from 3 mice per genotype) **F)** Confocal image and 3D reconstruction of K63-Ubiquitin^+^ foci (red) and nucleus (DAPI blue) and nuclear ubiquitin (yellow) in 18hr cultured MuSCs. Ubiquitin signal was defined as nuclear if the overlap ratio between Ubiquitin^+^ foci and nuclear volume was above 0.8. **G)** Confocal image and 3D reconstruction of PARKIN (red), SC35/SRRM2^+^ speckles (green) and nucleus (DAPI, blue) in freshly isolated wild-type MuSCs. **H)** Proportion of nuclear speckles labeled by PARKIN in freshly isolated wild type MuSCs. (*n*=64 cells from 6 mice)**I-J)** Quantification of total speckle content per nucleus and volume of individual speckles in *_MuSC_Park2^+/+^*and *_MuSC_Park2^-/-^* MuSCs. (*n*= 23-24 cells from 3 mice per genotype).**K)** Confocal image and 3D reconstruction of speckles (SC35/SRRM2, green) and nucleus (DAPI, blue) in freshly isolated *_MuSC_Park2^+/+^*and *_MuSC_Park2^-/-^* MuSCs. All data are presented as mean ± SEM. ns: not significant, *: p<0.05, **: p<0.01, ***: p<0.001, ****: p<0001 on unpaired two-tailed *t* tests or one-way ANOVAs comparing *_MuSC_Park2^+/+^* and *_MuSC_Park2^-/-^*.

Confocal microscopy and 3D reconstruction confirmed that while cytosolic PARKIN represented most of the cellular content (90-97%), a distinct nuclear fraction (2-10%) was consistently detected across all MuSC states (Fig. 5B-D, Fig S3A). Cytosolic PARKIN levels were highest in *in situ* fixed quiescent MuSCs and declined upon activation (Fig. 5C), consistent with changes in total PARKIN expression (Fig. 1G). In contrast, the nuclear PARKIN fraction increased as MuSCs transitioned from quiescence to activation and proliferation (Fig. 5D). Moreover, PARKIN loss reduced nuclear K63-ubiquitin-positive foci content (Fig. 5E-F), suggesting active participation of PARKIN in nuclear substrate ubiquitination and processes required for MuSC expansion.

Within the nucleus, PARKIN displayed a punctate staining pattern distributed across euchromatic regions, preferentially occupying the loose chromatin domains surrounding dense DAPI-positive heterochromatin foci. A fraction of PARKIN puncta showed partial juxtaposition with γH2AX-positive DNA damage foci^32^, though this association was infrequent (Fig. S3B-C). More frequently, PARKIN puncta were found in proximity to SC35/SRRM2-positive nuclear speckles, with approximately 6% of speckles displaying adjacent PARKIN labeling on snapshot images — a 3-fold higher frequency than observed at DNA damage foci (Fig. 5G-H). Notably, *_MuSC_Park2^-/-^* cells displayed significantly reduced content and size of SC35/SRRM2-positive foci (Fig. 5I-K).

Nuclear speckles, which form through Liquid-Liquid Phase Separation, serve as storage and assembly sites for spliceosomal components and splicing factors ^33,34^. Given the reduced expression of splicing-related genes in the DEG analysis (Fig. 5A and S2.3) and the altered speckle morphology in *Park2^-/-^*MuSCs (Fig. 5I-K), we investigated potential splicing defects through re-analysis of our RNA-seq data at the mRNA isoform level. Loss of PARKIN resulted in Differential Transcript Usage (DTU) affecting 636 genes, 90 of which also exhibited changes in overall gene expression levels (Fig. 6A). Among the 546 genes with DTU alone, approximately half showed isoform switches between different protein-coding transcripts (Fig. 6A-B). Critically, the remaining events represented hallmarks of impaired splicing fidelity: intron retention (26%), and non-coding processed transcripts (12%), nonsense-mediated decay (NMD)-inducing alterations (6%), collectively accounting for 44% of splicing alterations (Fig. 6B).

**Figure 6:**
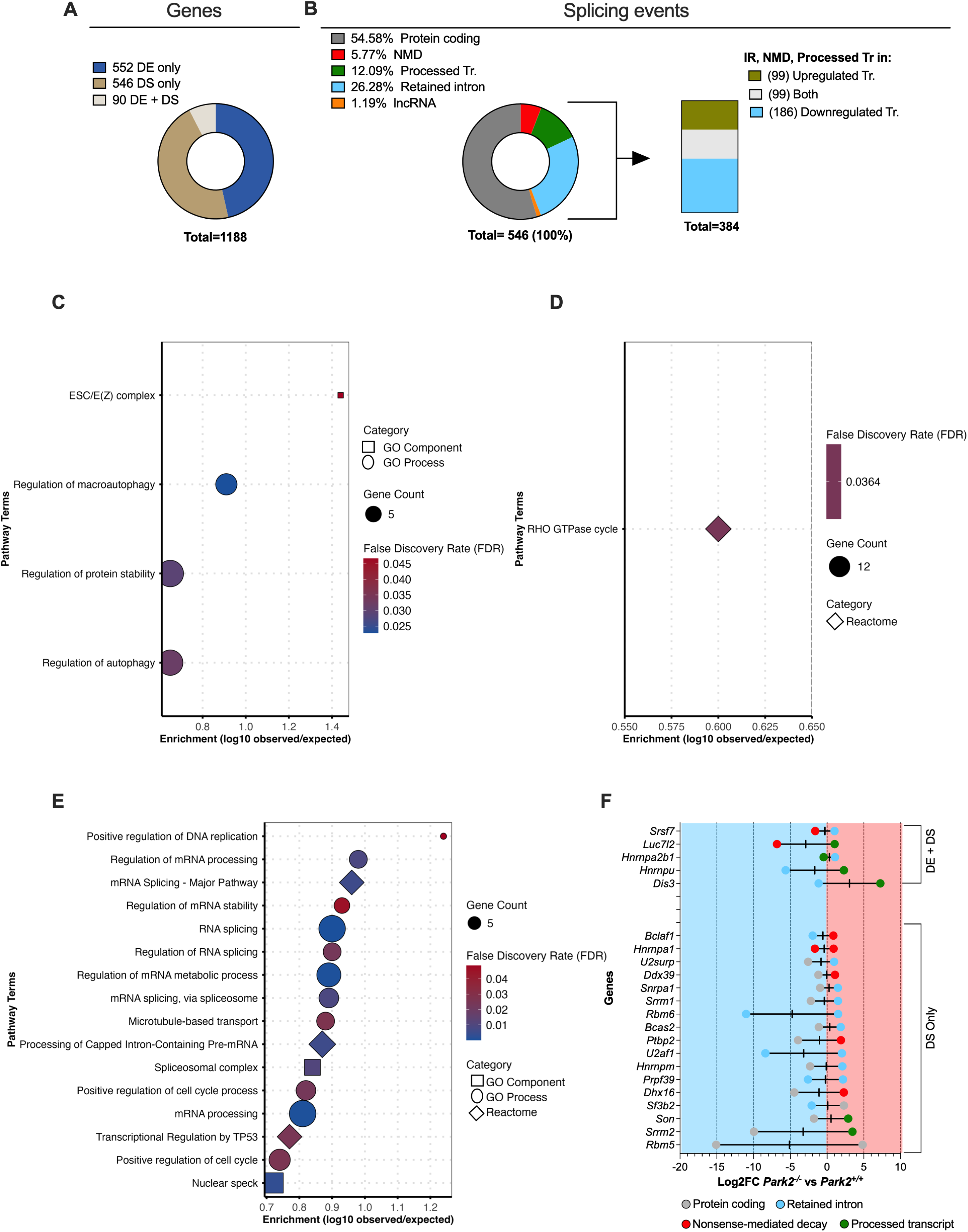
PARKIN deficiency induces self-targeting splicing defects that entrap spliceosomal machinery in non-productive isoforms. **A)** Transcriptomic landscape of PARKIN-deficient quiescent MuSCs. Left: Distribution of affected genes by type of alteration—differential expression only (DE only, blue), differential transcript usage only (DS only, tan), or both (DE+DS, white; total n=1,188 genes). **B)** Directionality of non-productive transcript changes. Among genes with IR, NMD, or processed transcript isoforms (n=384 events), stacked bar indicates the proportion where these non-productive isoforms are upregulated in *Park2*^-/-^ (green, n=99), downregulated in *Park2*^-/-^ (blue, n=186), or show bidirectional changes with both up- and downregulated non-productive isoforms (white, n=99). **C-E)** Gene Ontology enrichment analysis stratified by directionality of non-productive transcript changes. Bubble plots show enriched biological processes (GO:BP), cellular components (GO:CC), and Reactome pathways. **(C)** Enrichment analysis of genes with downregulated non-productive isoforms (n=186). **(D)** Enrichment analysis of genes with bidirectional changes showing both upregulated and downregulated non-productive isoforms (n=99). **(E)** Enrichment analysis of genes with upregulated non-productive isoforms (n=99), revealing selective enrichment for RNA splicing, mRNA processing, and spliceosomal machinery pathways. **F)** Isoform switching in core spliceosome and nuclear speckle genes. For 22 representative splicing machinery genes, log2 fold-change (*Park2*^-/-^ vs *Park2*^+/+^) of individual transcript isoforms grouped by whether genes show both differential expression and splicing (DE+DS, top) or splicing changes only (DS Only, bottom). The predominant pattern shows coordinated downregulation of protein-coding transcripts concurrent with upregulation of non-productive variants.

To understand the impact of these splicing changes, we examined pathway enrichment across different categories of affected genes. Genes with downregulated non-productive transcripts showed enrichment in broad cellular processes including macroautophagy and protein stability regulation, while genes exhibiting both upregulated and downregulated non-productive isoforms displayed diffuse enrichment patterns primarily related to cell cycle processes (Fig. 6C-D). In contrast, genes in which non-productive transcript isoforms were specifically upregulated in *Park2^-/-^* MuSCs revealed a striking and selective enrichment for splicing machinery components—including spliceosome constituents, RNA processing factors, and nuclear speckle proteins (Fig. 6E). Among these enriched genes, 22 core spliceosome and speckle components exhibited a characteristic pattern: loss of protein-coding variants coupled with accumulation of non-productive isoforms containing retained introns or destined for NMD (Fig. 6F). This selective pattern of aberrant splicing factor transcripts, coupled with the reduced SC35/SRRM2^+^ speckle integrity in *Park2^-/-^*MuSCs, points to dysregulated splicing machinery organization.

Intriguingly, the splicing defects in *Park2^-/-^* MuSCs showed overlap with intron retention programs characteristic of quiescent stem cells. Recent studies have demonstrated that quiescent muscle stem cells naturally exhibit widespread intron retention, which is resolved during activation in a tightly regulated manner ^15^. Comparison of genes with upregulated intron retaining variants in *Park2^-/-^* MuSCs tothose showing intron retention during deep quiescence (GSE113631 ^15^) identified 41 overlapping candidates (Fig. S4A). This overlap was significantly enriched for RNA splicing and mRNA processing pathways (Fig. S4B) and formed an interconnected network centered on core spliceosomal components (Fig. S4C). This molecular signature suggests that while *Park2^-/-^* MuSCs exhibit premature activation and commitment (Fig. 3F-I), they paradoxically retain key splicing machinery genes in an intron-retained, quiescence-like state, which could jeopardize proliferative expansion.

Given these abnormalities, we directly tested MuSC proliferative capacity. Dual EdU/Ki67 staining was performed on cultured EDL myofibers at 48 and 72 hours to track cell cycle progression. MuSC cluster size was significantly reduced in *_MuSC_Park2^-/-^* myofibers at both time points (Fig. 7A). At 48 hours, an increased proportion of non-proliferating cells (Ki67^-^/EdU^-^) was observed, suggesting delayed cell cycle re-entry (Fig. 7B-C). By 72 hours, *_MuSC_Park2^-/-^* cultures exhibited multiple cell cycle defects: reduced actively cycling cells (Ki67^+^/EdU^+^), increased G1/S phase delay (Ki67^+^/EdU^-^), and premature cell cycle exit (Ki67^-^/EdU^+^) (Fig. 7B-C). Interestingly, consistent with our previously published findings^13^, no difference in cluster size were observed in myofibers from *Pink1^-/-^* mice compared to their controls suggesting that the proliferation defects observed in PARKIN deficient MuSCs may be mechanistically distinct from the PINK1/PARKIN mitophagy pathway and the mitochondrial changes described earlier.

**Figure 7:**
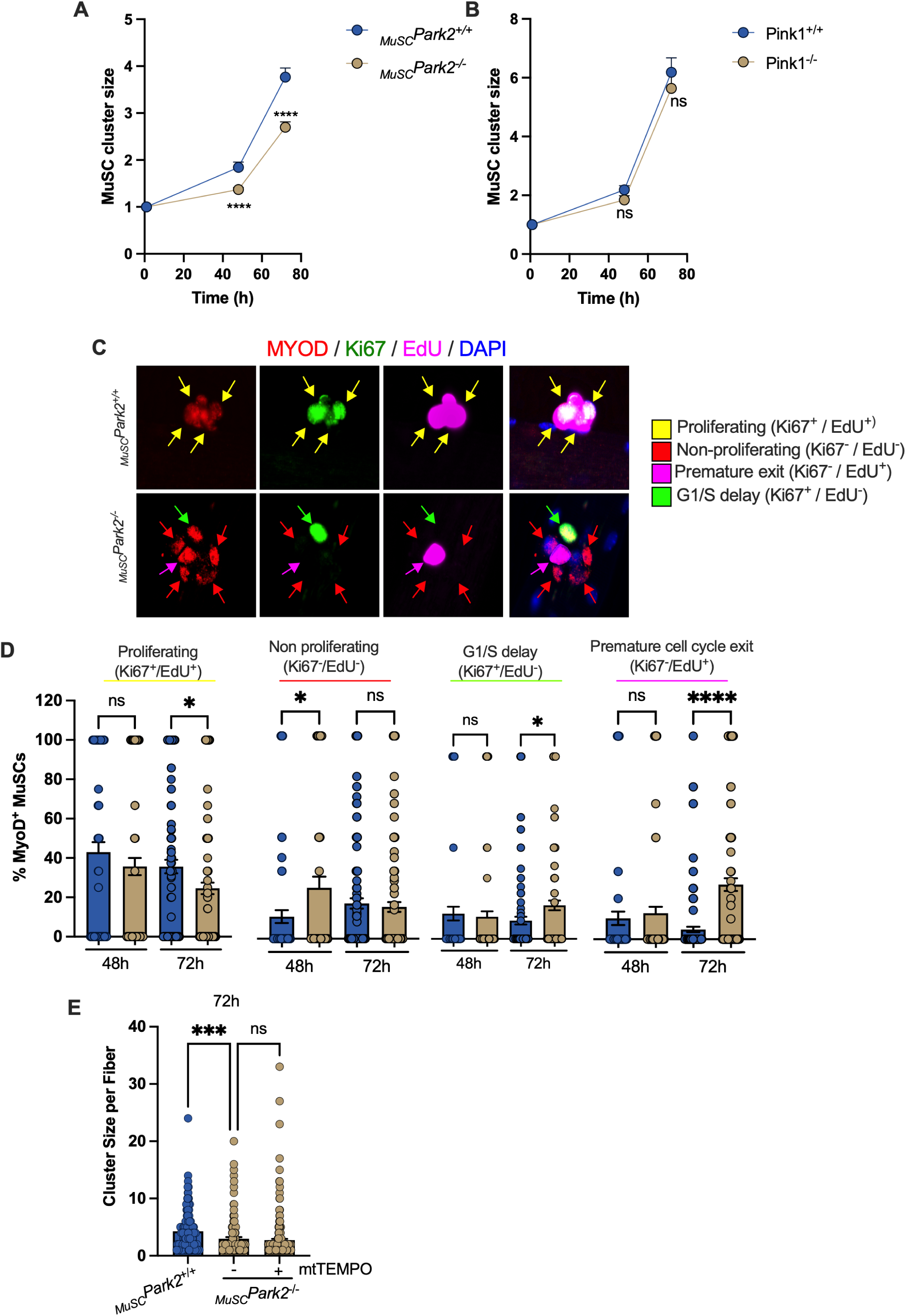
PARKIN is required for normal cell cycle regulation and expansion of the MuSC pool. **A)** Number of MuSCs per cluster in EDL fibers from *_MuSC_Park2^+/+^* and *_MuSC_Park2^-/-^* mice after 48 and 72h in culture (n=83-162 clusters from 4-7 fibers isolated from 3 mice per genotype). **B)** Number of MuSCs per cluster in EDL fibers from *Pink1^+/+^* and *Pink1^-/-^*mice after 48 and 72h in culture in the Cairns et al. study ^13^. **C)** Representative images of MYOD / Ki67 / EdU immunostainings in EDL fibers. Arrow colors illustrate the various cell cycle states (proliferating, non-proliferating, premature exit, G1/S delay). **D)** Proportion of MYOD^+^ MuSCs found in each the cell cycle state at 48 and 72h in culture. **E)** MuSC cluster size in EDL fibers cultured in absence or presence of 10 µM mitoTEMPO for 24h (n=52-135 MuSCs from 5 fibers from 4 mice per genotype). All data are presented as mean ± SEM. ns: not significant, *: p<0.05, **: p<0.01, ***: p<0.001, ****: p<0001 on one-way ANOVAs.

In this regard, while MitoTEMPO treatment effectively rescued fate specification defects in PARKIN-deficient MuSCs (Fig. 4E) as it did previously in PINK1-deficient MuSCs^13^, it failed to restore proliferative capacity (Fig. 7D). This dissociation between mitochondrial rescue and persistent cell cycle dysfunction suggests that PARKIN supports MuSC expansion through nuclear functions—potentially involving chromatin organization, splicing regulation, and mitotic progression—that operate independently of its canonical mitophagy role.

## 3 DISCUSSION

Mechanisms that couple mitochondrial state to nuclear reprogramming are fundamental to stem cell function yet remain poorly understood. Here, we identify PARKIN as a regulator of this coordination in muscle stem cells. PARKIN sustains mitophagy in quiescent MuSCs, preserves mitochondrial features that support balanced self-renewal and commitment decisions, and regulates the expression of nuclear genes encoding oxidative phosphorylation components. Unexpectedly, we uncovered a nuclear pool of PARKIN that localizes predominantly to interchromatin space, with focal association with nuclear speckles. Loss of PARKIN reduces K63-linked ubiquitination in the nucleus, disrupts speckle organization, induces splicing defects that lock splicing machinery genes in an intron-retained, quiescence-like state, and triggers aberrant activation signaling that limits proliferative expansion. Together, these findings reveal dual mitochondrial and nuclear roles for PARKIN that couple metabolic state to transcriptional reprogramming during stem cell state transitions and are essential for effective muscle regeneration.

### 3.1 Mitophagy: A gatekeeper of quiescence and regulator of fate decision

Mitochondria are now recognized as central regulators of stem cell fate, with their metabolic properties directly influencing the balance between self-renewal and differentiation ^5,9,12,13,31^. Quiescent stem cells, including MuSCs, maintain a limited mitochondrial network characterized by fatty acid oxidation, reduced oxidative phosphorylation, and low membrane potential and ROS emission—features that support quiescence while limiting oxidative damage ^28,29,35–37^. Exit from this metabolic state is required for activation and progression toward differentiation ^28,37,38^ underscoring the need for mechanisms that preserve mitochondrial properties during quiescence.

Our data identify mitophagy as a key mechanism underlying this preservation. Using *in situ* fixation to maintain native quiescence, we show that mitochondrial colocalization with autophagolysosomes and PARKIN recruitment are highest in truly quiescent MuSCs and markedly reduced in freshly isolated unfixed cells that undergo early activation during dissociation (Fig. 1). These findings demonstrate that propension to mitophagy is an intrinsic feature of quiescence rather than a consequence of isolation-induced stress. Upon activation, MuSCs rapidly suppress mitophagy, as evidenced by reduced *Parkin* expression, loss of PARKIN mitochondrial localization, and decreased mitochondrial engagement with autophagolysosomes (Fig. 2). Our genetic ablation study demonstrates that PARKIN is functionally important to sustain mitophagy consistent with our previous observations in germline PINK1-deficient mice ^13^. These results position the PINK1/PARKIN pathway as a gatekeeper of quiescence and a regulator of the metabolic transition from quiescence to activation.

Quiescent MuSCs, owing to their low metabolic activity, are not prone to mitochondrial damage from wear and tear. However, their characteristically low mitochondrial membrane potential^4,39,40^ —a well-established trigger for PARKIN recruitment ^19,41,42^ —naturally predisposes mitochondria toward PINK1/PARKIN-mediated mitophagy, functioning as a preventive mechanism to maintain organelles with uniformly low activity. When this quality control is defective, mitochondria escape surveillance, become polarized, and emit ROS that drives cells toward activation. Consistent with this model, PARKIN-deficient quiescent MuSCs displayed increased mitochondrial membrane potential, network fragmentation, and downregulation of nuclear-encoded OXPHOS genes—metabolic and transcriptional hallmarks of quiescence exit ^15,25,28,31,40^ (Fig. 4). Notably, upstream biogenesis regulators remained normally expressed (Fig. 4F), indicating these changes reflect altered cell state rather than direct PARKIN control of mitochondrial biogenesis. The same logic applies physiologically: natural MuSC activation leads to PARKIN suppression and rapid loss of mitochondrial colocalization (Fig. 1), concomitant with membrane polarization. Although mitochondrial ROS was not measured directly in PARKIN-deficient MuSCs, we previously reported elevated ROS in PINK1-deficient MuSCs ^13^, and scavenging mitochondrial ROS with MitoTEMPO fully restored self-renewal/commitment balance in PINK1 ^13^ and PARKIN-deficient cells (Fig. 5E). Collectively, these data support mitochondrial ROS as a key signaling intermediate linking defective PINK1/PARKIN-dependent preventive mitophagy to altered fate decisions.

### 3.2 A nuclear role for PARKIN in ubiquitination, splicing, and proliferative competence

Previous studies have reported nuclear localization of PARKIN in neurons ^23,43,44^, cardiac cells ^21^, and cancer models ^22^, typically under stress conditions (hypoxia ^21^, DNA damage ^32^), where PARKIN has been implicated in mitotic regulation through Cdc20/Cdh1 ^22^ and in transcriptional control, including ERRα-dependent programs during hypoxia ^21^. Our findings indicate that MuSCs harbor a constitutive pool of nuclear PARKIN under basal conditions that rises rapidly as cells transition from quiescence to activation, suggesting that nuclear PARKIN is not exclusively a response to exogenous stress but may be an integral feature of MuSC state transitions, although we cannot fully exclude that isolation-associated stimuli contribute to its nuclear accumulation. The mechanisms governing nuclear import of PARKIN remain incompletely understood, as PARKIN lacks a canonical nuclear localization sequence; however, post-translational modifications such as nitrosylation and SUMOylation have previously been shown to modulate the partitioning of PARKIN between the cytosol and nucleus^23^. We did not observe changes in ERRα targets ^21^, PPAR-related pathways ^21^, or PARIS (ZNF746) ^21^ as a result of PARKIN deficiency, arguing against the transcriptional roles attributed to nuclear PARKIN in other cell types as a primary mode of its activity in MuSCs.

Within the nucleus, PARKIN exhibited a punctate distribution predominantly across loose chromatin domains surrounding dense DAPI-positive heterochromatin, consistent with an euchromatic localization (Fig. 5B, 5G and S3). Within this interchromatin space, focal association with SC35/SRRM2-positive nuclear speckles was observed, though PARKIN was not exclusively confined to these structures. This subnuclear distribution is consistent with, but not limited to, a role in splicing regulation, and leaves open the possibility that nuclear PARKIN engages additional clients or nuclear processes, including transcriptional regulation, as has been proposed in other contexts^21,23^. Nuclear speckles act as reservoirs and assembly sites for splicing factors and undergo structural changes in response to shifts in transcription and RNA-processing demand ^33,34^. PARKIN-deficient MuSCs exhibited reduced speckle content and size (Fig. 5I-K), together with broad transcriptomic signatures consistent with splicing dysregulation, including intron retention and loss of productive isoforms among multiple splicing-related genes. Notably, several of these intron-retaining transcripts overlap with intron retention programs characteristic of quiescence ^15^. Our data thus suggest that PARKIN contributes to the transition from quiescence-associated RNA-processing programs to the splicing-competent state needed for effective proliferative expansion.

Although our data point to a role for nuclear PARKIN in maintaining context-appropriate splicing, they do not exclude contributions from other nuclear processes. It is noteworthy that our transcriptomic profiling was performed on freshly isolated MuSCs, a largely quiescent state with only very early transcriptional signs of activation ^15,25^. At this stage, mitotic programs have not yet been engaged ^25^, making it more likely that the downregulation of pathways related to mitosis and chromosome organization observed in *Park2^-/-^*MuSCs reflects downstream consequences, whereas early splicing abnormalities represent a more proximal nuclear defect. Critically, our experimental design does not allow us to formally dissect the respective contributions of nuclear versus mitochondrial/cytosolic PARKIN to the observed phenotypes. It is nonetheless noteworthy that proliferative defects were not observed in PINK1-deficient MuSCs (Fig. 7B and ^13^) despite comparable mitophagy impairment, and that MitoTEMPO failed to restore proliferative capacity (Fig. 7E) despite rescuing fate specification (Fig. 4E). These observations raise the possibility that proliferative competence may be particularly sensitive to the nuclear functions of PARKIN, though this interpretation remains speculative in the absence of tools that selectively perturb each pool independently.

How PARKIN functions in quiescent MuSCs, where splicing output is low, remains unresolved. However, our isoform-level analyses revealed that numerous spliceosome and nuclear-speckle genes exhibit isoform switching toward non-productive biotypes in *Park2^-/-^*MuSCs, including variants with retained introns, nonsense-mediated-decay potential, or processed-transcript annotations. Among these, SON and SRRM2 are particularly notable: together they constitute the core scaffold essential for nuclear-speckle assembly and for concentrating splicing factors within these condensates ^45,46^. Notably, ubiquitin-dependent pathways, such as those involving the speckle-associated deubiquitinase USP42, are known to modulate LLPS behavior and nuclear-speckle organization ^47,48^, indicating that ubiquitination can influence condensate dynamics within the nucleus. Consistent with this transcript-level disruption, SC35/SRRM2 staining appeared more diffuse and less abundant in *Park2^-/-^* MuSCs, indicating altered speckle organization at the protein level. Because nuclear speckles promote early spliceosome engagement and efficient co-transcriptional splicing of associated genes ^33,34,45^, the shift of SON, SRRM2, and other splicing regulators toward non-productive isoforms provides a plausible explanation for the broader transcriptomic signatures of splicing dysregulation and reduced speckle volume observed in *Park2^-/-^* MuSCs. Although we did not undertake a mechanistic dissection of nuclear PARKIN, these transcriptomic and imaging signatures are consistent with a model in which PARKIN helps sustain a splicing-competent nuclear-condensate environment during the earliest stages of activation. Future studies will be required to determine whether this reflects direct ubiquitin-dependent regulation of speckle-resident proteins or an indirect consequence of impaired splicing of key scaffold factors.

Overall, our results indicate that PARKIN supports MuSC activation and expansion through dual metabolic and nuclear influences. Mitochondrial PARKIN preserves a quiescence-compatible mitochondrial state that maintains balanced fate decisions, while the combined transcriptomic and imaging signatures in *Park2^-/-^* MuSCs, including isoform switching in core speckle components and altered SC35/SRRM2 organization, suggest that PARKIN also contributes to sustaining a nuclear RNA-processing environment permissive for productive proliferation.

## 4 METHODS

### 4.1 Animal care

#### Methods

*Park2^loxP/loxP^* mice were obtained from Lexicon Pharmaceuticals ^49^ and bred with *Pax7CreERT2^GAKA^*mice obtained from Jackson’s laboratory (*B6.Cg-Pax7^tm1(cre/ERT2)Gaka^/J*) to develop a Tamoxifen inducible and conditional knockout of *PARKIN* in PAX7^+^ adult MuSCs. *Park2^loxP/loxP^*-*Pax7Cre^+/0^*were used to knockout *Parkin* in MuSCs while *Park2^wt/wt^- Pax7Cre^+/0^* were used as controls. At 7-9 weeks of age, all mice received Tamoxifen (500 mg/kg) (Sigma T5648, 50 mg/mL dissolved in corn oil) administration via oral gavage for five consecutive days, followed by a seven-day washout period before any subsequent procedures were carried out. CAG-RFP-EGFP-LC3 reporter mice (C57BL/6-Tg(CAG-RFP/EGFP/Map1lc3b)1Hill/J, Jackson Laboratory) were used to examine lysosomal activity in MuSCs. Mice were bred according to the recommended hemizygote × noncarrier breeding strategy. All experiments on animals were approved by the University of Ottawa Institutional Animal Care Committee and conducted according to the directives of the Canadian Council on Animal Care. Mice were maintained in ventilated cage racks by groups of 3-5 mice. All mice of both sexes were kept on a regular 12–12 hr light-dark cycle and had access to food and water *ad libitum.* Animals were used at 8-12 weeks of age for experiments and euthanized by cervical dislocation.

### 4.2 Cardiotoxin injury

#### Methods

Cardiotoxin was administered to the TA in accordance with previously published protocol ^28^.The uninjured contralateral TA was used as control in all experiments. Muscle was prepared and sectioned as described in^13,28^.

### 4.3 Muscle tissue preparation

#### Methods

Cardiotoxin was administered to the TA in accordance with the protocol described by Baker *et al.*, 2022. ^28^ The un-injected contralateral TA was used as control in all experiments. Muscle was prepared and sectioned as described in ^13,28^.

### 4.4 Single EDL fiber isolation and culture

#### Methods

EDL muscles were harvested and digested in collagenase B. Single fibers were isolated by gentle trituration with wash media and transferred to culture media for 24-72 hours, with or without EDU (2 hours) or MitoTempo (24 hours).

### 4.5 Muscle stem cell purification and culture

#### Methods

Muscle stem cells were isolated from hindlimb muscle of experimental mice. On ice, muscle was rapidly dissected (within 5 minutes of sacrifice) and placed immediately in a mixture of collagenase B and dispase II at 37°C. After digestion, muscle slurry was purified as described previously^13^, and labelled with (1) phycoerythrin-conjugated SCA-1, CD45, CD31, and CD11b, and (2) 647-conjugated α integrin-7 and VCAM-1. MuSC were then sorted by FACS using MoFlo XDP cell sorter.^13^

*In situ* fixed MuSCs were isolated according to ^25^ with slight modifications. Muscles were dissected on ice and placed immediately into ice cold 0.5% PFA where they were chopped into 1 mm pieces and incubated for one hour. Fixed muscles were washed and digested in collagenase-dispase for 1.5 hours at 37°C with a GentleMACS Octo dissociator. The muscle slurry was then purified with a series of 800g spins and serial filtering to remove debris. The cleared muscle slurry was then stained as described above, but with GFP conjugated CD34 replacing VCAM-1.

### 4.6 Primary myoblast isolation and culture

#### Methods

Muscle stem cells were isolated by FACS and plated at 3000 cells/cm^2^ on Matrigel coated dishes. Myoblast culture was then expanded and passaged on Matrigel (P1-2) or gelatine (P3+) coated plates. Primary myoblasts were cultured in growth media containing Ham’s 10 nutrient mix supplemented with 20% FBS (v/v), 1% penicillin/streptomycin (P/S) (v/v), 2.5ng/L bFGF and 2% Ultroser-G (v/v). Following the first passage, growth media was switched to UG-omitted media ^50^.

### 4.7 Hematoxylin & Eosin staining

#### Methods

Flash frozen muscle was cut, stained and imaged as described previously ^13^. Fiber cross sectional area (CSA) was determined on three 1000 µm^2^ regions of interest (ROI) using Fiji (ImageJ) ^13^.

### 4.8 Immunofluorescence

#### 4.8.1 Muscle sections

##### Methods

Antigen retrieval was performed on TA muscle sections, followed by blocking and incubation with primary antibodies against PAX7, MYOD, MYOG, and secondary fluorescent antibodies. These sections were imaged with EVOS Fl Auto2 microscope (20x). MuSCs were counted in 500 μm^2^ ROIs and normalized to number of fibers present in each ROI^13^.

#### 4.8.2 Fate determination in EDL fibers

##### Methods

Fibers were permeabilized and stained as described previously ^13^, then imaged with EVOS FL Auto2 at 100X, or Leica Thunder Imager at 40X. Images from Leica Thunder then underwent computational clearing prior to analysis. Fate decision determination was performed by counting number of PAX7/MYOD/MYOG expressing cells per cluster. Cluster size was counted as all DAPI positive cells in a cluster. To measure proliferative capacity, fibers were treated with EDU for 2 hours prior to fixation. EDU detection was performed with BaseClick EDU detection kit (647). These fibers were then incubated with primary antibodies to PAX7/MYOD and Ki67.

#### 4.8.3 Purified MuSCs and cultured myoblasts

##### Methods

Following fixation and permeabilization, cells were blocked and incubated with primary antibodies against TOM20, LC3, PARKIN, γH2AX, SC35/SRRM2 orK63-Ubiquitin, followed by secondary antibodies and DAPI. Cells were washed and imaged with Zeiss LSM880 AxioObserver Z1 (63x). Cells imaged by Zeiss LSM880 AxioObserver Z1 underwent deconvolution by AiryScan.

### 4.9 Confocal image processing and quantification of mitophagy

#### Methods

FACS-purified MuSCs labeled with DAPI, TOM20, LC3, PARKIN, γH2AX, SC35/SRRM2 and K63-Ubiquitin, were imaged and processed using the Airyscan function. Volume reconstruction and surface/surface colocalization analysis was performed in Imaris using fixed settings.

### 4.10 RNA isolation and qPCR

#### Methods

To perform quantitative real-time qPCR, RNA was isolated from freshly purified MuSCs or cultured myoblasts using PicoPure RNA isolation kit. RNA was then reverse transcribed into cDNA using iScript Reverse Transcription Supermix for RT-qPCR and amplified using SsoAdvanced Universal SYBR Green Supermix. qPCR was run against housekeeping genes HPRT or GAPDH.

### 4.11 Transcriptomic profiling and differential isoform analysis

#### Methods

1000-3000 MuSCs were sorted directly into lysis buffer and immediately underwent cDNA synthesis following lysis using Smart-Seq HT kit according to the protocol described in^51^. cDNA libraries were prepared using Nextera XT DNA Library Preparation Kit, size selected, then indexed with Nextera XT Index Kit. Sequencing was performed on Illumina NovaSeq 6000 at the SickKids Center for Applied Genomics. Reads acquired were paired-end 105bp. Quality control was assessed using FastQC.

Raw RNA-sequencing reads were aligned to the mouse reference genome (GRCm39) using STAR^52^, and transcript-level abundances were quantified using RSEM^53^. Differential gene expression (DGE) analysis was performed in R using edgeR^54^ with RUVseq normalization to account for batch effects. Differential transcript usage (DTU) was assessed using the diffSplice function in edgeR and the DTUrtle^55^ package to identify isoform switching events independent of total gene expression changes. Functional interpretation was performed using Over-Representation Analysis (Gprofiler^56^) and Gene Set Enrichment Analysis (GSEA^57^) with standard and custom gene sets, and protein-protein interaction networks were constructed using the STRING database^58^. Detailed parameters and statistical thresholds are provided in Supplementary Methods.

### 4.12 ATP Assay

#### Methods

ATP content was determined in freshly sorted MuSCs using the Cell-Titre Glo luminescence viability assay kit. The contribution of mitochondria to ATP provision was determined by comparing ATP levels in presence and absence of the ATP synthase inhibitor Oligomycin as per^28^.

### 4.13 Mitochondrial Membrane Potential

#### Methods

MuSCs were loaded with the potentiometric probe TMRE (4 nM, 30 minutes, 37°C), and immediately analyzed by flow cytometry on a BD LSR Fortessa flow cytometer. Cells treated with CCCP (5 µM) were used to confirm loss of fluorescence in response to uncoupling. Analysis of flow cytometry data was performed on FloJo 10.10.0.

### 4.14 Data mining

#### Methods

Publicly available transcriptomics datasets (GSE103162, GSE70736, GSE55490, GSE47177) ^12,17,25,27^ comparing quiescent and activated MuSCs were used for targeted analyses. For each dataset, the list of differentially expressed genes (q value<0.05) was extracted and used for targeted analyses of specific gene sets.

### 4.15 Statistics and reproducibility

For experiments performed in single FACS purified MuSCs, cultured EDL myofibers or myoblasts, a minimum of three different isolations/culture with a minimum of 10 MuSCs or fibers, or 3-10 distinct culture wells was analyzed. For experiments involving *in vivo* CTX injury, a minimum of 3-4 mice per genotype were used and for each TA muscle analyzed 3 distinct ROIs of 1000 µm^2^ each was included to cover a large proportion of the entire muscle. Values are reported as mean ± SEM and represented as bar graphs along with individual datapoints when appropriate. Unpaired two-sided t-tests with Welch’s correction were used to determine statistical difference when two means were compared. To compare more than two means, Brown-Forsythe and Welch ANOVA tests were used with Dunnett’s T3 multiple comparisons test performed for each pair of means compared. Outlier analyses were performed using the ROUT method with a 5% Q value cutoff. All statistical analyses were performed using GraphPad Prism 10.4.2.

### 4.16 Resource availability

#### Lead contact

Yan Burelle.

#### Material availability

Mouse lines used are available commercially at Jackson’s laboratory or can be obtained through Lexicon pharmaceuticals.

#### Source data

Raw RNAseq data is available on GEO (GSEXXXXX). List of differentially expressed and differentially transcribed genes and data used to generate figures are provided as supplemental data. All other data are available from the corresponding authors upon reasonable request.

## 6 AUTHOR CONTRIBUTIONS

Conception of hypothesis: YB. Design of the work: YB, MK, GC, MA, MG. Acquisition and analysis of the data: MG, MA, GC, MTM, AW, HL, NL. Drafting the manuscript: YB, MK, MG, GC. All authors: Analysis and interpretation of the data and approval of the final version of the manuscript.

## 8 GEOLOCATION INFORMATION

Ottawa, Ontario, Canada

## 9 ACKNOWLEDGEMENTS

The authors gratefully acknowledge services provided by the Louise Pelletier histology core (RRID: SCR_021737) and the Cell Biology and Image Acquisition Core (RRID: SCR_021845) funded by the University of Ottawa, the Natural Sciences and engineering Research Council of Canada, and the Canada Foundation for Innovation. The authors also acknowledge the expert contribution of Fernando Ortiz and Shahriar Sheikholeslami from Flow Cytometry & Cell Sorting Facility at OHRI for its contribution on FACS, RRID:SCR_023349 and Gareth Palidwor from the OHRI Bioinformatics Core.

## SUPPLEMENTALS

1. **Extended Methods**
2. **Source data for Figures**
3. **Supplementary Table 1- Gene level statistics:** This table reports gene level statistics including padj, logFC, and gene symbol.
4. **Supplementary Table 2- Integrated differential transcript utilization and differential expression analysis in *Park2^−/−^* vs *Park2^+/+^* MuSCs.** This table reports, for each gene, statistics from differential transcript utilization (DTU) and differential gene expression (DE) analyses, including FDR values, transcript-level log2 fold changes, and the identity and metrics of the most differentially used transcripts. Genes are categorized as undergoing DTU only, DE only, or both

## 12 EXTENDED METHODS

### 12.1 Key resources table

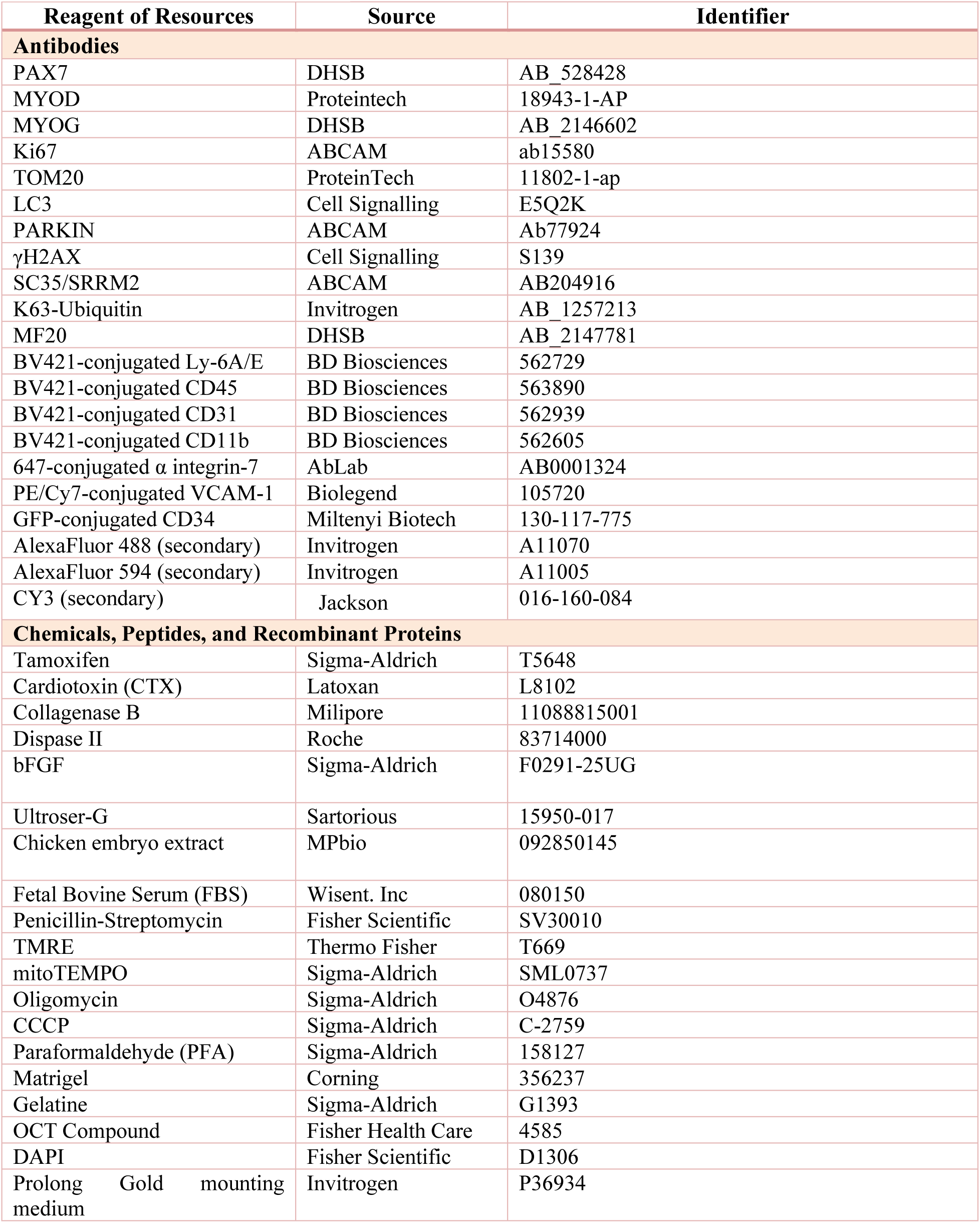
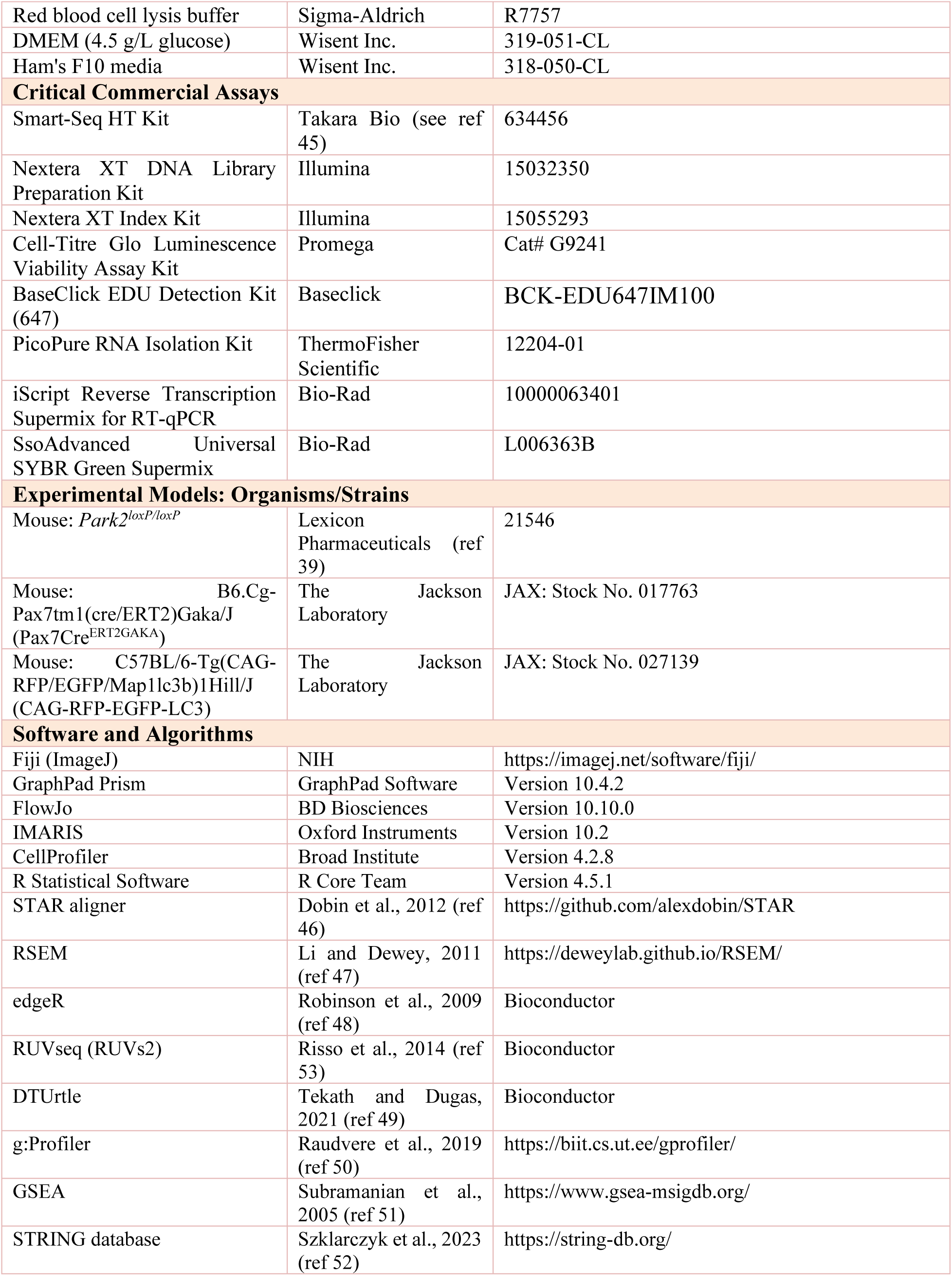
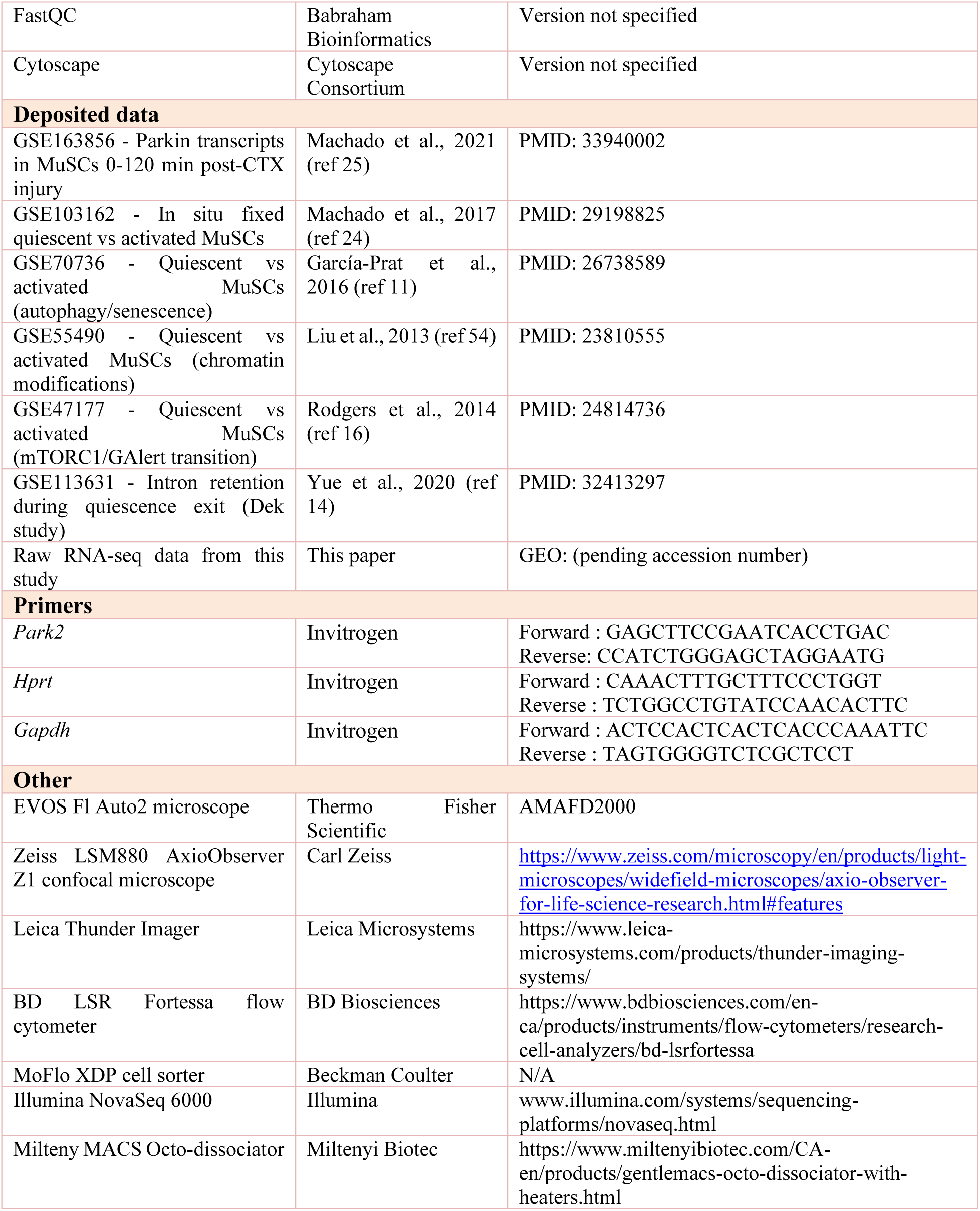

### 12.2 Animal care

*Park2^loxP/loxP^* mice were obtained from Lexicon Pharmaceuticals^49^ and bred with *Pax7CreERT2^GAKA^*mice obtained from Jackson’s laboratory (*B6.Cg-Pax7^tm1(cre/ERT2)Gaka^/J*) to develop a Tamoxifen inducible and conditional knockout of *PARKIN* in PAX7^+^ adult MuSCs. *Park2^loxP/loxP^*-*Pax7Cre^+/0^* were used to knockout *Parkin* in MuSCs while *Park2^wt/wt^- Pax7Cre^+/0^*were used as controls. At 7-9 weeks of age, all mice received Tamoxifen (500 mg/kg) (Sigma T5648, 50 mg/mL dissolved in corn oil) administration via oral gavage for five consecutive days, followed by a seven-day washout period before any subsequent procedures were carried out. CAG-RFP-EGFP-LC3 reporter mice (C57BL/6-Tg(CAG-RFP/EGFP/Map1lc3b)1Hill/J, Jackson Laboratory) were used to examine lysosomal activity in MuSCs. Mice were bred according to the recommended hemizygote × noncarrier breeding strategy. All experiments on animals were approved by the University of Ottawa Institutional Animal Care Committee and conducted according to the directives of the Canadian Council on Animal Care. Mice were maintained in ventilated cage racks by groups of 3-5 mice. All mice of both sexes were kept on a regular 12–12 hr light-dark cycle and had access to food and water *ad libitum.* Animals were used at 8-12 weeks of age for experiments and euthanized by cervical dislocation.

### 12.3 Cardiotoxin injury

Mice were injected subcutaneously with Buprenorphine (0.1 mg/Kg) and anesthetized by gas inhalation 30 min prior to Cardiotoxin (CTX) injection. CTX diluted in PBS (50 µL of 10 µM) was injected into the *Tibialis Anterior* (TA) muscle and the uninjured contralateral muscle was used as control. Mice recovered in a cage with a heating pad and were monitored for 24 hours post-CTX injection.

### 12.4 Muscle tissue preparation

At selected time points following CTX injection, TA muscles were dissected, weighed, and cut in half in a cross-sectional orientation. One portion was embedded in Optimal Cutting Temperature (OCT) compound and flash frozen in cooled isopentane, while the other portion was dropped in freshly prepared paraformaldehyde (2% w/v, 4 °C) and fixed for 30 minutes. Following fixation, TA muscles were washed twice with PBS for 5 minutes, twice with glycine (0.25M in PBS) for 10 minutes, and then left in 5% (w/v) sucrose for 2 hours, followed by 20% sucrose (w/v) for 2-3 days. TA muscles were then embedded in OCT compound, frozen in cooled isopentane and stored at −80°C. Tissues were cryo-sectioned in a cross-sectional orientation at a thickness of 14 µm and glass slides were stored at −80°C until further processing.

### 12.5 Single EDL fiber isolation and culture

The extensor digitorum longus (EDL) muscles were harvested immediately following cervical dislocation and digested in 0.5% (w/v) collagenase B for approximately 30-40 minutes at 37°C. Muscles were transferred in wash media (DMEM containing 4.5 g/L glucose and 1% (v/v) penicillin-streptomycin), and single fibers were obtained by gentle trituration of muscles with a wide bore pipette. Single fibers were then freed from debris through two successive washes. Following a 10 min rest period, fibers were either fixed in warmed PFA (2% w/v) for 10 min to capture the quiescent state or, some fibers were cultured over 12-72h (in DMEM 4.5 g/L glucose, 20% (v/v) FBS, 1% (v/v) chicken embryo extract, 1% (v/v) penicillin-streptomycin, 7.5 ng/mL βFGF) prior to fixation in order to track MuSC activation and commitment. Some fibers were treated with EDU (2 hour) or MitoTempo (24 hours). Fixed fibers were kept at 4 °C until further processing.

### 12.6 Muscle stem cell purification and culture

Hindlimb muscles were harvested, pooled, and digested in 1% (w/v) Collagenase-B and 0.4% (w/v) Dispase II in Hams-F10 media, using the Milteny MACS Octo-dissociator SLICE_FACS program for 27 minutes as per^28^. Following digestion, the muscle slurry was filtered (100µm mesh size) and centrifuged (10 min, 400xg). The cell pellet was resuspended in red blood cell lysis buffer and incubated for 5 minutes to remove residual erythrocytes. Cells we then spun down (5 min, 400xg) and washed in FACS buffer (5mM EDTA, 10% (v/v) FBS in 1xPBS) before being resuspended in 1 mL of FACS buffer containing the following antibodies: A) PE-conjugated Ly-6A/E, CD45, CD31 and CD11b to remove non-MuSCs present in the population, and B) 647-conjugated α-integrin-7 and VCAM-1 to positively select α-integrin-7 ^+^/VCAM^+^ MuSCs. Following incubation with antibodies (30-45 min at 4°C), the volume was topped to 15 mL with FACS buffer and cells were centrifuged 5 min at 400g. The resulting pellet was resuspended in 1 mL of FACS buffer, filtered through a 50 µm CellTrics filter into a 5 mL round bottom tube. Cells were then sorted by FACS at the Flow Cytometry facility at the Ottawa Hospital Research Institute (OHRI).

For *in situ* fixed MuSCs, muscles were dissected on ice and placed immediately into ice cold 0.5% PFA where they were chopped into 1 mm pieces and incubated for one hour. Following digestion, the muscle slurry was filtered (100µm mesh size) and centrifuged (10 min, 800xg) to remove debris. This step is repeated with a 50µm mesh size. The cleared muscle slurry was then stained as described above, but with GFP conjugated CD34 replacing VCAM-1.

### 12.7 Primary myoblast isolation and culture

Muscle stem cells were isolated by FACS and plated at 3000 cells/cm^2^ on Matrigel coated dishes. Myoblast culture was then expanded and passaged on Matrigel (P1-2) or gelatine (P3+) coated plates. Primary myoblasts were cultured in growth media containing Ham’s 10 nutrient mix supplemented with 20% FBS (v/v), 1% penicillin/streptomycin (P/S) (v/v), 2.5ng/L bFGF and 2% Ultroser-G (v/v). ^48^

### 12.8 Hematoxylin & Eosin staining

Cross sections from flash frozen samples were stained with Hematoxylin and Eosin (H&E) at the uOttawa Histology Core Facility. Images were taken at 20x magnification on an EVOS Fl Auto2 microscope and digitally stitched to reconstitute entire muscle sections. For each muscle, myofiber numbers and minimum Ferret diameter were measured on three 1000 µm^2^ regions of interest (ROI) using Fiji (ImageJ). Myofiber cross sectional area (CSA) was calculated using the equation CSA = π x (minimum Ferret diameter/2)^13^.

### 12.9 Immunofluorescence

#### 12.9.1 Muscle sections

Tissue sections were submitted to antigen retrieval in citrate buffer (10 mM sodium citrate, 0.05% Tween-20, pH=6.0), followed by 1 hour blocking in 3% (w/v) BSA. Slides were then incubated overnight at 4°C with primary antibodies against PAX7. Slides were then washed 5 times with 1xPBS, and fluorescence-conjugated secondary antibodies, (AlexaFluoro 488, AlexaFluoro 594 or CY3) diluted in 3% BSA were added to sections. Following incubation at room temperature for 60 mins, sections were washed 5 times and slides were mounted using Prolong Gold with DAPI. Images were taken at 20x (0.4 NA) magnification on an EVOS Fl Auto2 microscope, digitally stitched to reconstitute entire muscle sections, and analyzed using Fiji (ImageJ). For each muscle section, the number of MuSCs was counted in three distinct 500 µm^2^ ROIs and normalized by the number of muscle fibers present in each ROI^13^.

#### 12.9.2 Fate determination in EDL fibers

Single EDL myofibers were permeabilized in 0.1% (v/v) Triton-X and 100 mM glycine, followed by blocking in blocking solution (5% (v/v) horse serum, 2% (w/v) BSA and 0.1% (v/v) Triton-X in PBS) for 5 hours at room temperature. EDU detection was performed with BaseClick EDU detection kit. Myofibers were incubated overnight at 4°C with a combination of the following primary antibodies diluted in blocking solution: PAX7, MYOD, Ki67 and MYOG. Following incubation with secondary antibodies (AlexaFluoro 488, AlexaFluoro 594) for 1 hour at room temperature, myofibers were mounted on charged glass slides using Prolong Gold and imaged either using a 100x/1.28 PL Apo Oil immersion on an EVOS Fl Auto2 microscope or a 40×/0.95 HC PL Apo CORR-DRY objective on a Leica Thunder microscope. Each myofiber was manually inspected throughout its thickness and along its full length to capture the entire population of MuSC.

#### 12.9.3 Purified MuSCs and cultured myoblasts

Cells were fixed in 4% PFA, washed 3X in PBS and quenched in 50mM NH_4_Cl. Cells were permeabilized in 0.1% (v/v) Triton X-100 and washed with PBS. Cells were blocked in 10% (v/v) FBS for 1 hour and incubated with primary antibodies in 5% FBS overnight. Primary antibodies included TOM20, LC3, PARKIN, γH2AX, SC35/SRRM2 and K63-Ubiquitin. Cells were incubated with secondary antibodies (AlexaFluoro 488, AlexaFluoro 594), then incubated with DAPI 300nM for 5 minutes. Cells were then washed, cover slipped, and imaged either on an a Zeiss LSM880 AxioObserver Z1 (63X magnification 1.4 NA, Oil, Plan-Apo).

### 12.10 Confocal image processing and quantification of mitophagy

FACS purified MuSCs labeled with DAPI, TOM20, LC3, PARKIN, γH2AX, SC35/SRRM2 and K63-Ubiquitin were imaged on a Zeiss LSM880 confocal microscope and processed using the Airyscan function. To control batch effects and sampling biases, cells from *MuSCPark2^-/-^*and *MuSCPark2^+/+^* mice were always isolated and processed in parallel for each experimental day, and multiple cells were randomly imaged in each experiment. To control for image processing and quantification biases, analyses were batch processed in the IMARIS software using fixed settings for the intensity and voxel size threshold values, and surface/surface colocalization parameters. Analyses were performed on MuSCs from uninjured mice that were fixed immediately after FACS purification or 4h after spontaneous *in vitro* activation. In some experiments, muscle was fixed in situ prior to MuSC purification in order to capture the native quiescent state.

### 12.11 Transcriptomic profiling and differential isoform analysis

1000-3000 MuSCs were sorted directly into lysis buffer and immediately underwent cDNA synthesis following lysis using Smart-Seq HT kit according to the protocol described in ^51^. cDNA libraries were prepared using Nextera XT DNA Library Preparation Kit, size selected, then indexed with Nextera XT Index Kit. Sequencing was performed on Illumina NovaSeq 6000 at the SickKids Center for Applied Genomics.

To characterize the molecular landscape and isoform dynamics in stem cell populations, we employed a high-resolution RNA-sequencing pipeline integrating genomic alignment, transcript quantification, and differential utilization modeling. Raw sequencing reads were subjected to adapter trimming and quality filtering to remove low-confidence bases and technical artifacts before alignment to the mouse reference genome (GRCm39) and transcriptome using STAR (Spliced Transcripts Alignment to a Reference) with parameters optimized for accurate mapping of multi-isoform genes and novel splice junctions^52^.

Transcript-level abundances were quantified using RSEM ^53^ (RNA-Seq by Expectation-Maximization), which employs an Expectation-Maximization algorithm to provide maximum likelihood estimates of transcript expected counts, effectively resolving mapping ambiguities among highly similar isoforms characteristic of stem cell pluripotency factors^53^. Gene-level counts were derived by summarizing expected counts across all constituent transcripts. Statistical analysis of differential gene expression (DGE) was performed in R using the edgeR package^54^, with RUVseq (RUVs2)^60^ applied to normalize data and account for technical variation and batch effects across stem cell cultures. DGE was calculated using the glmQLFit function on gene-summarized expected counts, with significant genes identified by empirical Bayes quasi-likelihood F-tests. To resolve transcript-level complexity, we employed two complementary approaches: the diffSplice function in edgeR to detect variations in relative transcript expression within genes, and the DTUrtle package^55^ (run_drimseq function) applied to RSEM-quantified transcript proportions to model differential transcript usage (DTU) and identify functional isoform switching events occurring independently of total gene expression changes.

Functional interpretation of differentially expressed genes was performed using Over-Representation Analysis (Gprofiler package^56^) and Gene Set Enrichment Analysis (GSEA package^57^) in R, with all genes detected in the sequencing run used as the background environment. For Gprofiler, the g:SCS (shortest common superstring) algorithm was used for multiple hypothesis testing correction with a default alpha threshold of 0.05, and only GO terms annotating 300 genes or fewer were considered to filter out large annotations with limited interpretative value. For GSEA, genes were ranked by signed p-value (sign of fold change × - log₁₀(p-value)) to preserve both magnitude and direction of expression changes, gene sets smaller than 1500 genes were analyzed, and significance was determined by normalized enrichment score (NES) >1.5 and adjusted p-value <0.1. Both standard and custom gene sets representing focused pathways were employed to identify enrichments. High-confidence protein-protein interaction networks and their associated enrichment terms were identified using the STRING database^58^.

### 12.12 ATP Assay

Freshly isolated MuSCs were divided in triplicates of 10 000 cells per well in 96-well transparent F bottom microplates. Luminescence was detected using Cell-Titre Glo luminescence viability assay kit (Promega, G9241), and cellular ATP content was determined by means of a standard curve. To determine mitochondrial dependent ATP production, cells were treated with or without the mitochondrial ATP synthase inhibitor Oligomycin (20 µM) for 30 minutes prior to detection.

### 12.13 Data mining

Publicly available transcriptomics datasets (GSE103162, GSE70376 GSE55490, GSE47177) ^12,17,25,27^ comparing quiescent and activated MuSCs were used for targeted analyses. For each dataset, the list of differentially expressed genes (q value<0.05) was extracted and used for targeted analyses of specific gene sets.

### 12.14 Statistics and reproducibility

For experiments performed in single FACS purified MuSCs, cultured EDL myofibers or myoblasts, a minimum of three different isolations/culture with a minimum of 10 MuSCs or fibers, or 3-10 distinct culture wells was analyzed. For experiments involving *in vivo* CTX injury, a minimum of 3-4 mice per genotype were used and for each TA muscle analyzed 3 distinct ROIs of 1000 µm^2^ each was included to cover a large proportion of the entire muscle. Values are reported as mean ± sem and represented as bar graphs along with individual datapoints when appropriate. Unpaired two-sided t-tests with Welch’s correction were used to determine statistical difference when two means were compared. To compare more than two means, Brown-Forsythe and Welch ANOVA tests were used with Dunnett’s T3 multiple comparisons test performed for each pair of means compared. Outlier analyses were performed using the ROUT method with a 5% Q value cutoff. All statistical analyses were performed using GraphPad Prism 10.4.2.

## 13 SUPPLEMENTARY FIGURES

**Figure S1: Data supplement for Figure 1 and 2. A-B)** Total volume of mitochondria and LC3^+^ structures per cell quantified following reconstruction of *in situ* fixed, freshly sorted quiescent and 4h *in vitro* activated wild type MuSCs (*n*=25-36 cells from 3 mice in each group). **C)** Representative confocal images of autophagolysosomes in unfixed MuSCs from EGFP-RFP-LC3 reporter mice at the indicated time points. High green/red ratio indicates a predominance of neutral pH compartments with poorly acidified inactive lysosomes while low green/red ratio indicates a predominance of acidified mor active lysosomes. **D)** Proportion of MuSCs harboring neutral (inactive) vs acidified (active) autophagolysosmes quantified from the total volume of GFP^+^/RFP^+^ and RFP^+^ only puncta per cell (n=3-4 separate experiments). **E-F)** Total volume of mitochondria and LC3^+^ structures per cell quantified following reconstruction of freshly sorted quiescent and 4h *in vitro* activated MuSCs from *_MuSC_Park2^+/+^* and *_MuSC_Park2^-/-^* mice (*n*=41-52 cells from 3 mice in each group). All data are presented as mean ± SEM. *: p<0.05, **: p<0.01 on one-way ANOVAs.

**Figure S2: Data supplement for Figure 4-5. A)** STING network of significantly downregulated mitochondrial proteins (log2FC < 0, adjusted p value < 0.05) in *_MuSC_Park2^-/-^* vs *_MuSC_Park2^+/+^* MuSCs. Proteins related to oxidative phosphorylation and mitochondrial translation are highlighted in color (complex I: green, complex III: blue, complex IV: magenta, complex V: peach, mitochondrial ribosomal proteins: light purple). **B)** Gene set enrichment analysis (GSEA) enrichment map showing enriched Gene Ontology (GO) gene sets in *_MuSC_Park2^-/-^* vs *_MuSC_Park2^+/+^* MuSCs. GSEA was performed using a ranked gene list based on p value × sign(logFC). Only gene sets containing at least 10 genes and fewer than 1,500 genes were included. Significantly enriched gene sets were defined by an absolute normalized enrichment score (NES) > 1.5 and an adjusted p value < 0.001. The table on the right summarizes major pathway clusters identified based on gene set similarity scores. **C-F)** Leading edge focussed heatmaps of selected gene sets in *_MuSC_Park2^-/-^* vs *_MuSC_Park2^+/+^* MuSCs. On large heatmaps, gene symbols are omitted for clarity and to visually represent general patterns of change.

**Figure S3: Data supplement to Figure 5. A)** Confocal microscopy and 3D surface rendering of freshly isolated muscle stem cells (MuSCs) from wild-type mice, stained for PARKIN (red) and DAPI (blue). Top and bottom rows show XY and ZY orthogonal views, respectively. Left panels show the merged DAPI/PARKIN signal. Middle panels display cytoplasmic PARKIN (red) and nuclear PARKIN (yellow) reconstructions overlaid on the DAPI nuclear surface (blue mesh). Right panels show nuclear PARKIN foci (yellow) alone, cytosolic surfaces removed for clarity, confirming that PARKIN-positive structures reside within the nuclear volume. PARKIN foci were classified as nuclear when their overlap ratio with the DAPI-defined nuclear mask exceeded 0.8, and as cytoplasmic when this ratio was between 0 and 0.4. Scale bars, 5 µm. **B)** Confocal image and 3D reconstruction of PARKIN (red), γH2AX^+^ DNA damage foci (green) and nucleus (DAPI, blue) in isolated fixed type MuSCs. **C)** Proportion of DNA damage foci labeled by PARKIN in isolated fixed wild type MuSCs. (*n*=38 cells from 4 mice)

**Figure S4: Data supplement to Figure 6. A)** Overlap between genes with upregulated intron retaining transcripts in PARKIN-deficient MuSCs and genes with intron retention during deep quiescence (IRS>1000 in fixed QSC vs activated SC; GSE113631, ^15^. Venn diagram shows 41 overlapping genes. **B)** Pathway enrichment of the 41 overlapping genes demonstrates selective enrichment for RNA splicing and mRNA processing functions. **C)** Protein-protein interaction network of the 41 overlapping genes. Network generated using STRING database with high-confidence interactions (score >0.7). Red/orange nodes indicate genes annotated as spliceosomal or nuclear speckle components; gray nodes indicate other interacting proteins. Network demonstrates that genes trapped in intron retaining splicing states upon PARKIN loss form a highly connected cluster centered on core splicing machinery.

