## Supplementary figures and images for "Loss of Parkin Disrupts Nuclear and Mitochondrial Programs Required for Muscle Regeneration"

### Supplemental Figures

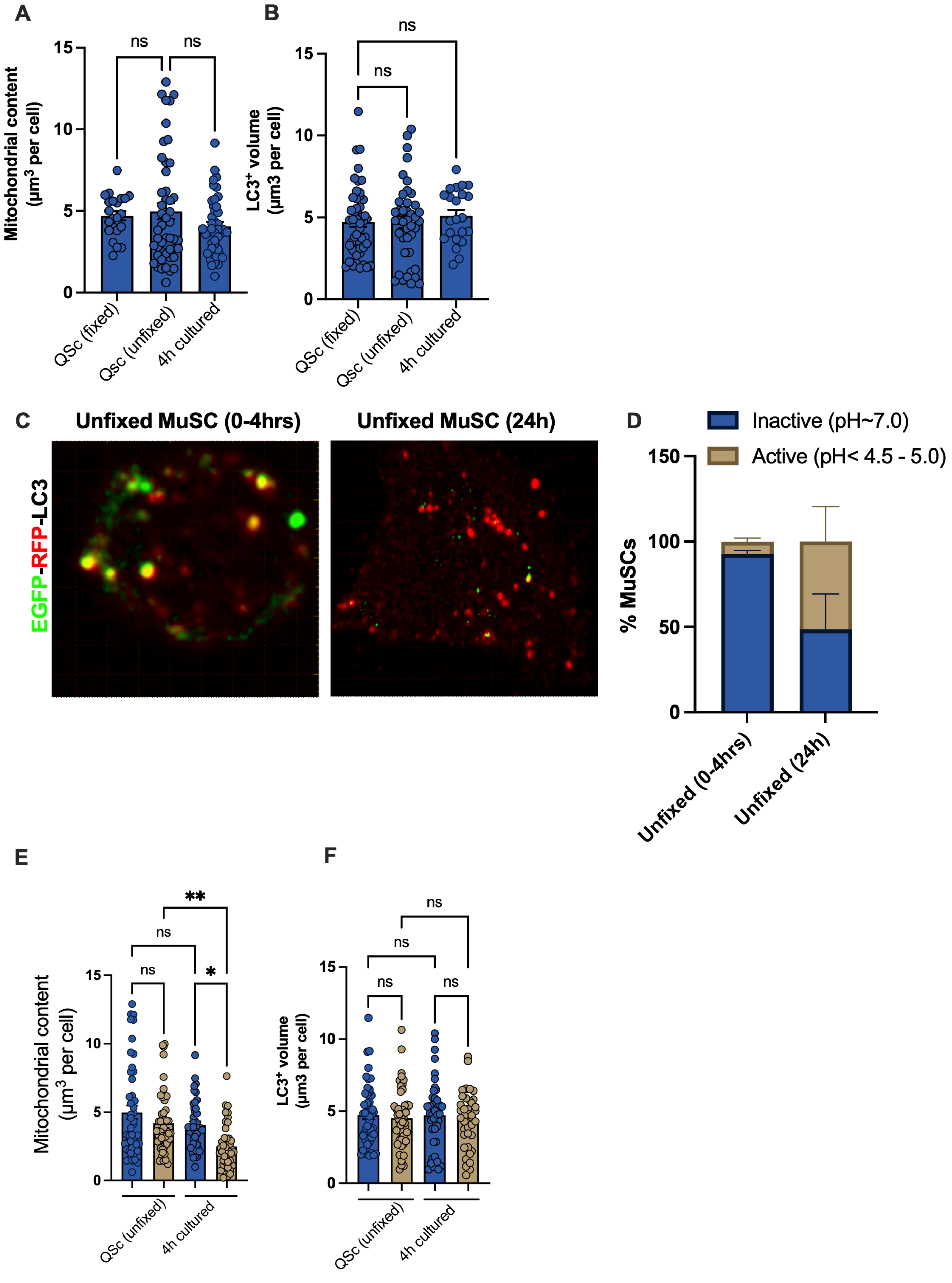

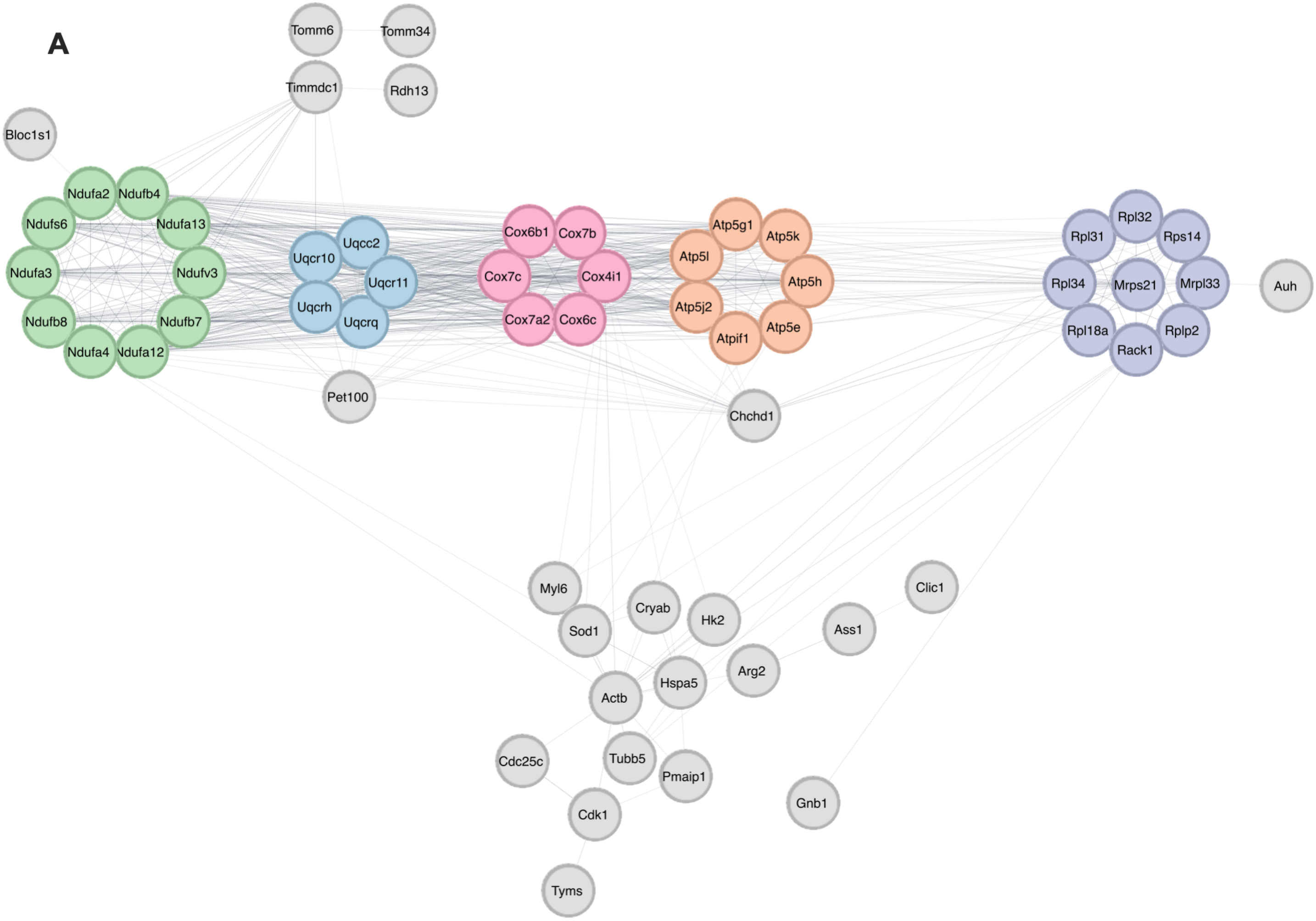

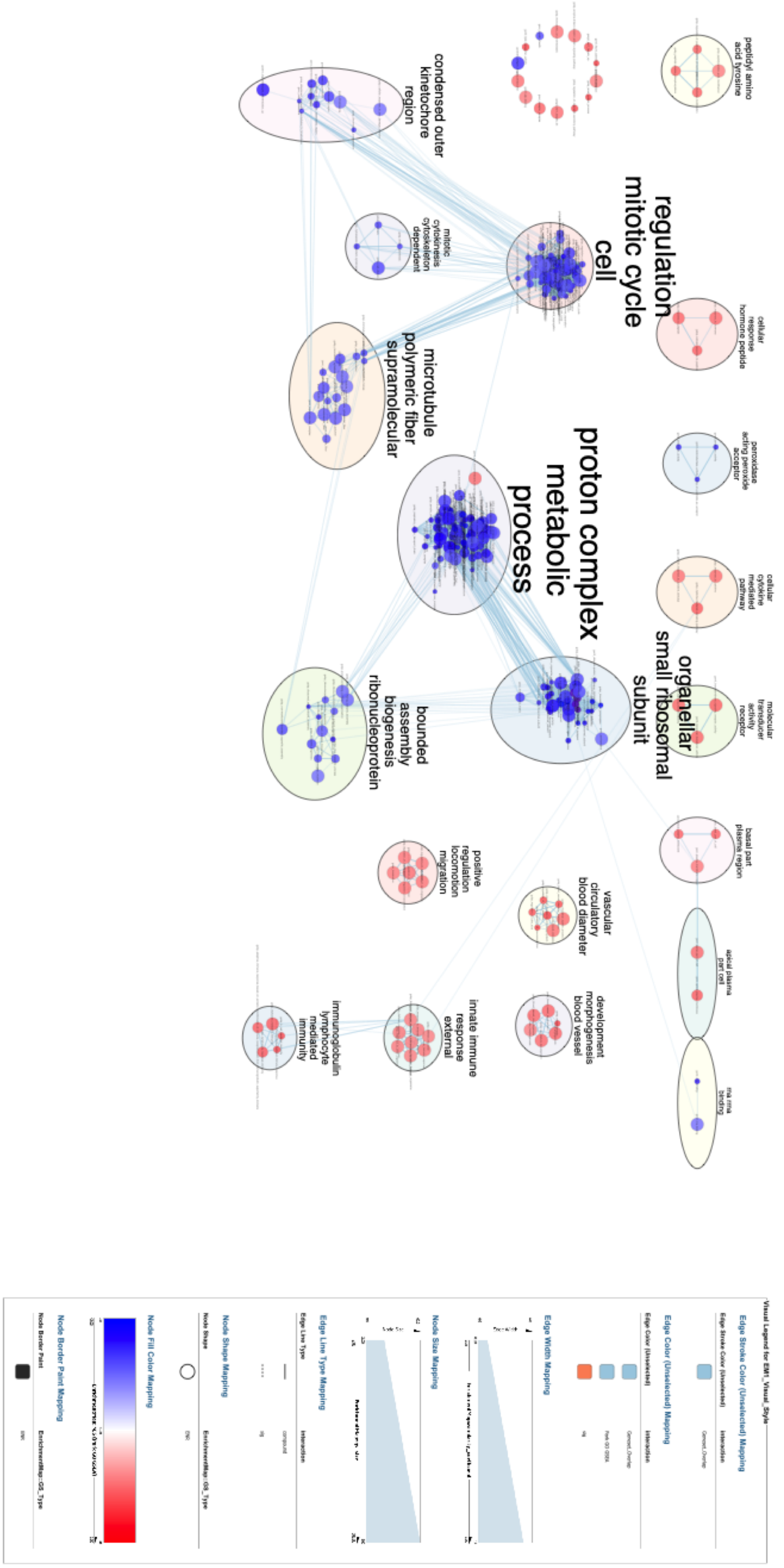

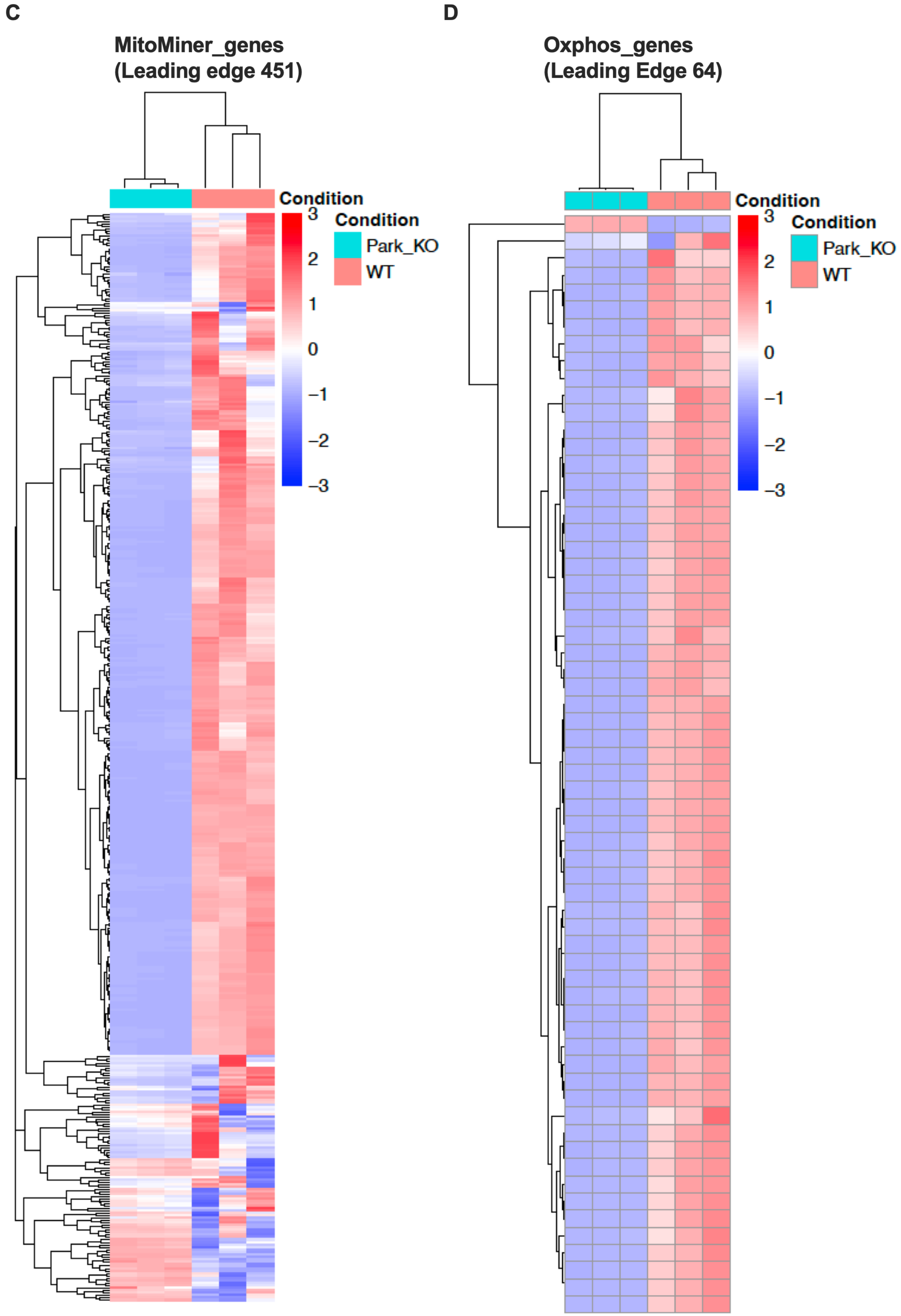

**E**

**Spliceosome (Leading edge 16)**

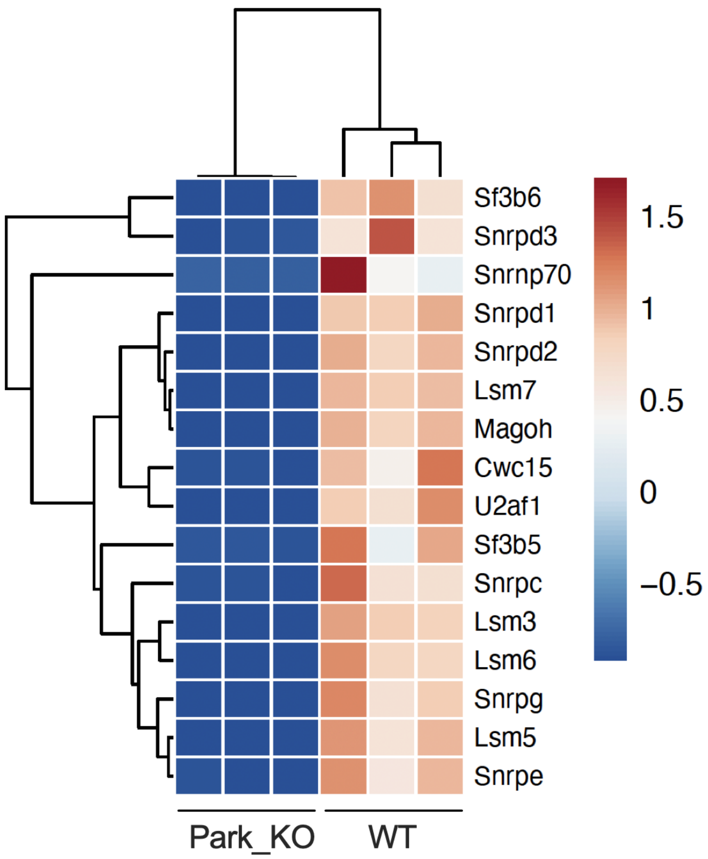


**F**

**Cell_cycle (Leading Edge 16)**

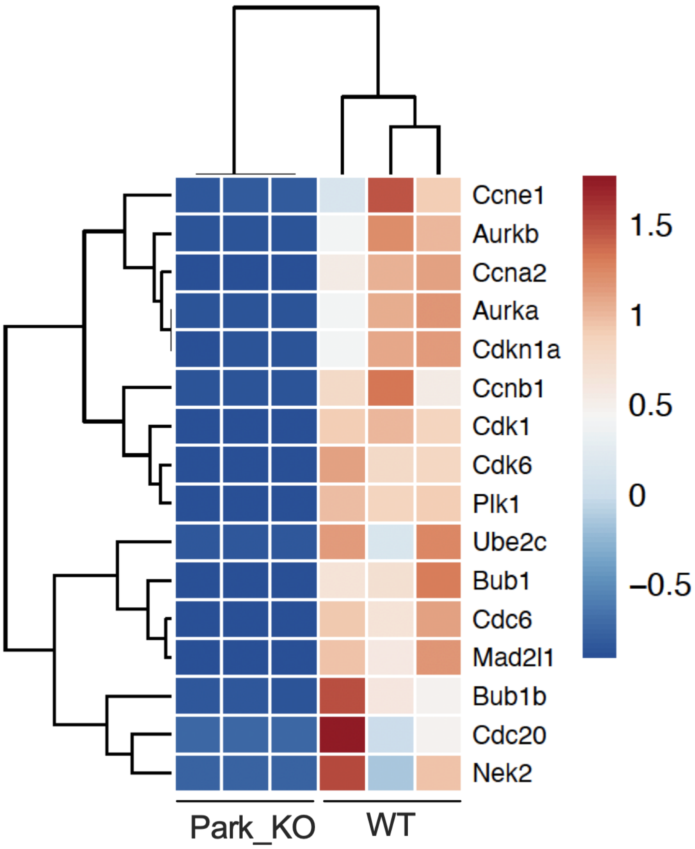


Figure S2

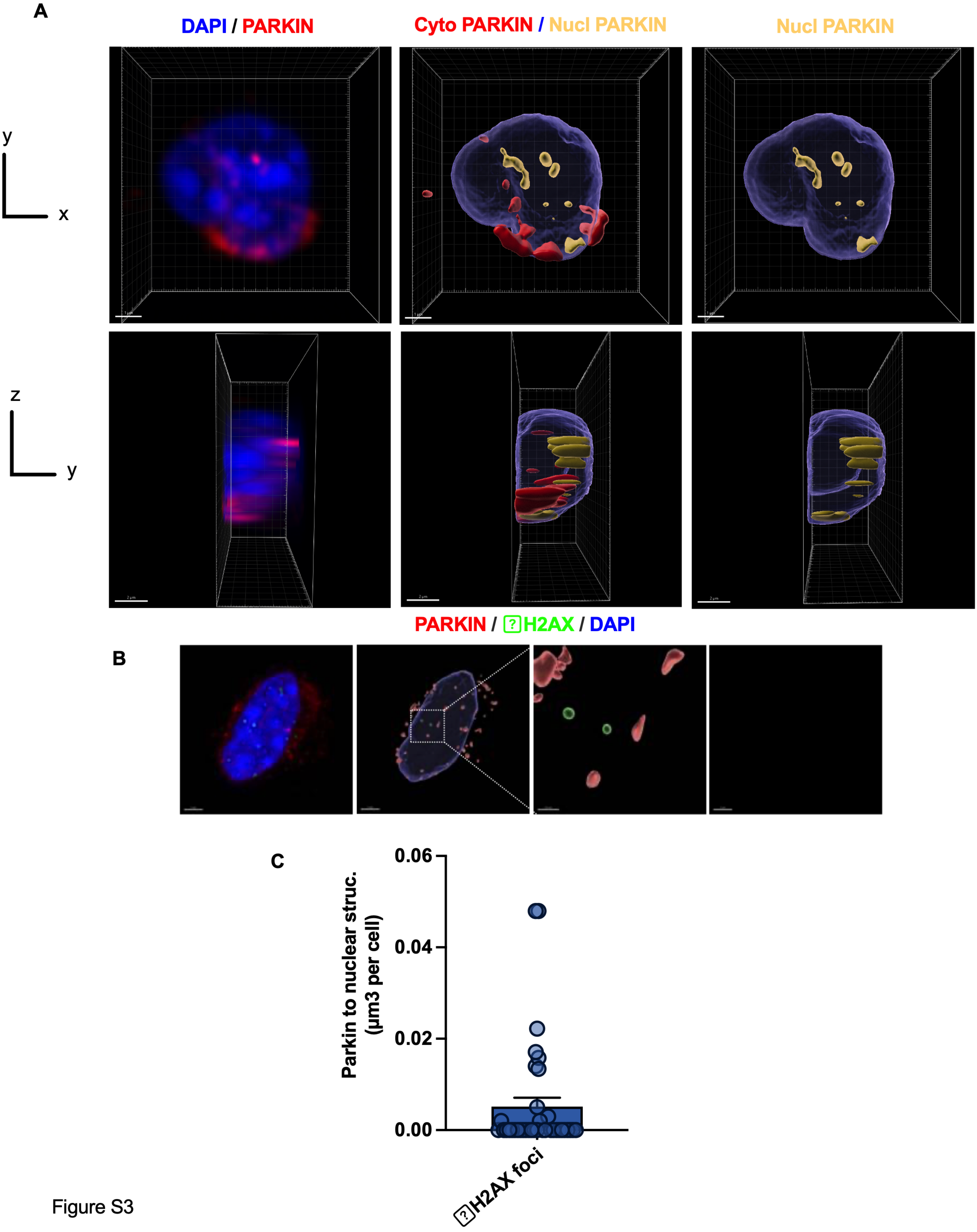

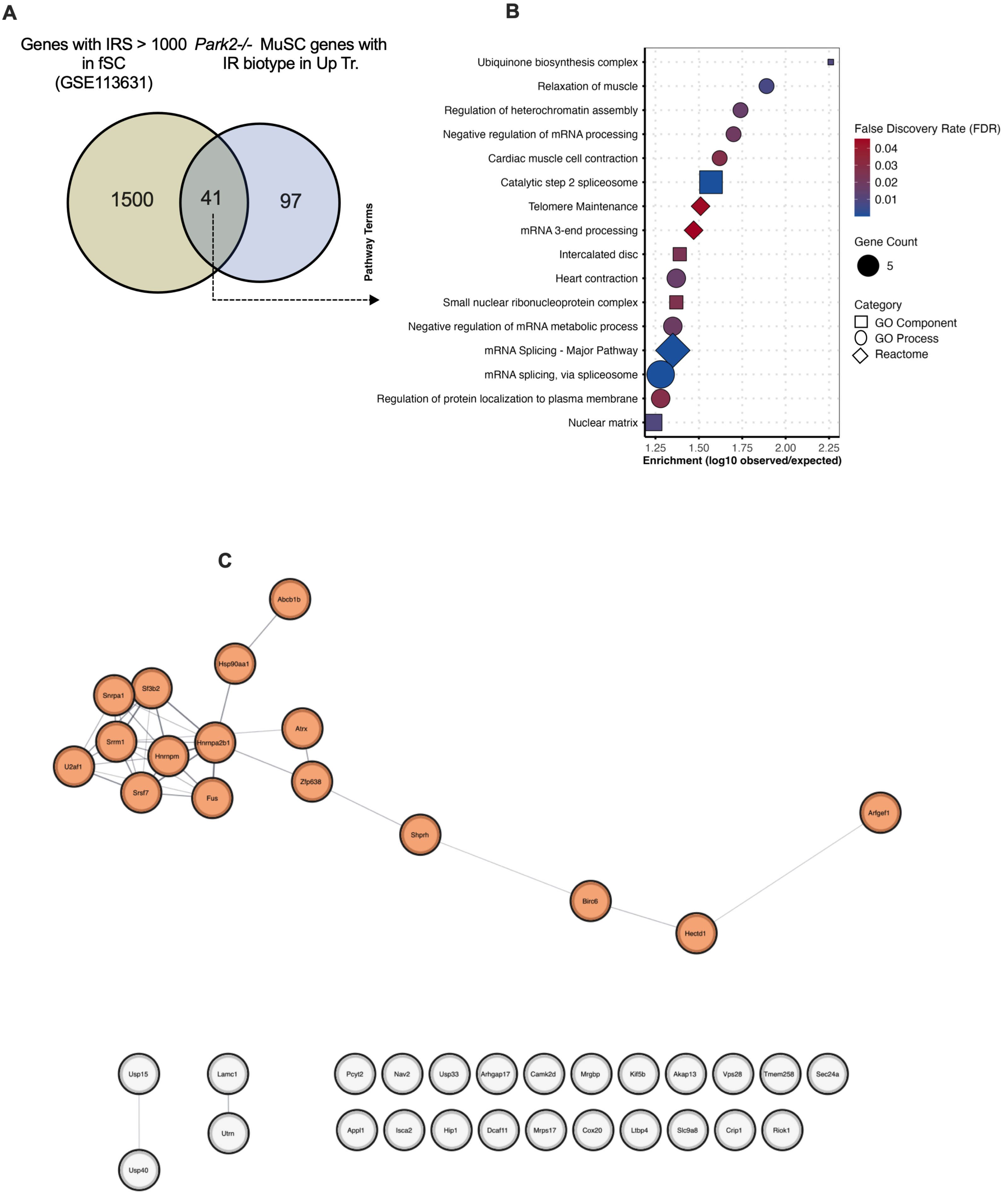
